# Physiological insight into the conserved properties of *Caenorhabditis elegans* acid-sensing DEG/ENaCs

**DOI:** 10.1101/2022.04.12.488049

**Authors:** Eva Kaulich, Patrick T. N. McCubbin, William R. Schafer, Denise S. Walker

## Abstract

Acid sensing ion channels (ASICs) are members of the diverse family of degenerin/epithelial sodium channels (DEG/ENaCs). They perform a wide range of physiological roles in healthy organisms, including in gut function and synaptic transmission, but also play important roles in disease, as acidosis is a hallmark of painful inflammatory and ischaemic conditions. We performed a screen for acid-sensitivity on all 30 subunits of the *C. elegans* DEG/ENaC family using Two-Electrode Voltage Clamp (TEVC) in *Xenopus* oocytes. We found two groups of acid-sensing DEG/ENaCs characterised by being inhibited or activated by increasing proton concentrations. Three of these acid-sensitive *C. elegans* DEG/ENaCs were activated by acidic pH, making them functionally similar to the vertebrate ASICs. We also identified four new members of the acid-inhibited DEG/ENaC group, giving a total of seven additional acid-sensitive channels. We observed sensitivity to the anti-hypertensive drug amiloride as well as modulation by the trace element zinc. Acid-sensitive DEG/ENaCs were found to be expressed in both neurons and non-neuronal tissue, highlighting the likely functional diversity of these channels. Our findings provide a framework to exploit the *C. elegans* channels as models to study the function of these acid-sensing channels *in vivo*, as well as to study them as potential targets for anti-helminthic drugs.

## Introduction

Acidosis can occur under healthy physiological conditions, such as during synaptic transmission (Du *et al*., 2014), as well as being a hallmark of a wide range of pathologies. Cells monitor tissue acidosis through membrane proteins including acid-sensing ion channels (ASICs) (Vina *et al*., 2013; Ortega-Ramirez *et al*., 2017). ASICs belong to the conserved family of Degenerin/Epithelial Sodium Channels (DEG/ENaC), non-voltage gated cation channels that are involved in a diverse range of cellular processes. As the name indicates, the family also includes mammalian ENaCs and *C. elegans* degenerins, as well as *Drosophila* pickpockets (PPK) and an array of representatives from across animal phyla. Electrophysiological approaches, particularly using *Xenopus* oocytes, have played an essential role in establishing the physiology of channel properties of this diverse family (Canessa *et al*., 1993; O’Brodovich *et al*., 1993; Canessa *et al*., 1994; Schild *et al*., 1997; Zhang & Canessa, 2002; Li *et al*., 2009).

Acid-sensing DEG/ENaC members across species can be classified into two groups, those activated and those inhibited by high proton concentrations. The former group includes the mammalian ASICs (Waldmann *et al*., 1997; Zhang & Canessa, 2002), zebrafish zASICs (Chen *et al*., 2007), human ENaCs (Collier & Snyder, 2009) and *Drosophila* PPK1 (Boiko *et al*., 2012). Vertebrate ASICs are closed at neutral pH and generate proton-activated inward currents, which increase with decreasing extracellular pH. However, the precise properties depend on the subunit composition, with the half activation pH varying from around 6.5 to 4.5 (Waldmann *et al*., 1997; Zhang & Canessa, 2002; Chen *et al*., 2007). Together they thus cover a significant pH range, with relevance to a diverse array of biological processes and contexts. Heteromeric human αβγENaC channels, on the other hand, are open at neutral pH with a maximal current at pH 6 and minimal current at pH 8.5 (Collier & Snyder, 2009), correlating well with the pH range in the collecting duct of the kidney and other epithelia where ENaCs are expressed. Rat αβγENaC currents are not altered over the same pH range, highlighting variation between species (Collier & Snyder, 2009). The second group of proton-sensitive DEG/ENaC members display inward currents at neutral pH, in the absence of additional stimulus, and are blocked by acidic pH. This group consisted of only three members, mouse ASIC5 (also called brain liver intestine Na^+^ channel (BLINaC)) (Wiemuth & Grunder, 2010), *Trichoplax* TadNaC6 (Elkhatib *et al*., 2019) and *C. elegans* ACD-1 (Wang *et al*., 2008; Wang *et al*., 2012), to which we have added two more *C. elegans* channels, ACD-5 and the heteromeric FLR-1/ACD-3/DEL-5 channel (Kaulich *et al*., 2021).

The evidence linking ASICs to important roles in neuronal health and disease makes them of particular interest, and tractable genetic model systems like *C. elegans* can facilitate insights into these processes. For instance, gain-of-function mutations in the *C. elegans* DEG-1, MEC-4 and UNC-8 subunits cause neuronal degeneration (Chalfie & Sulston, 1981; Bianchi *et al*., 2004; Wang *et al*., 2013). Introducing one of the causative mutations into human ASIC2 also causes neuronal cell death, suggesting that ASIC2 might also be involved in neurodegeneration (Waldmann *et al*., 1996). This hypothesis has received attention since ASIC2 is upregulated in patients with Multiple Sclerosis (MS), an inflammatory neurodegenerative disease and pharmacological blocking of ASICs can lessen clinical symptoms of inflammation and neuronal degeneration (Friese *et al*., 2007; Fazia *et al*., 2019). ASICs (and *C. elegans* DEG/ENaCs) are also targets of diverse non-steroidal anti-inflammatory drugs (NSAIDs) (Voilley *et al*., 2001; Voilley, 2004; Fechner *et al*., 2021), highlighting their importance as targets for treating pain and inflammation (Dulai *et al*., 2021). Finally, ASICs could offer potential as targets for anti-helminthic drugs. A variety of compounds is currently used to treat or prevent parasitic nematode infections, including imidazothiazoles and macrocyclic lactones, which target nicotinic acetylcholine receptors and glutamate-gated chloride channels, respectively (Wolstenholme & Rogers, 2005; Keiser & Utzinger, 2008; Williamson *et al*., 2009). Widespread resistance is a critical threat to both agriculture and human health, so effective alternatives are urgently needed.

*C. elegans* DEG/ENaCs are vastly expanded compared to their vertebrate members, with 30 subunit-encoding genes (compared to 4-5 in vertebrates). As the phylogram and the Sequence Similarity Network (SSN) in Figure 1 and Figure 2 show, they form distinct homology groups, with the vertebrate ENaCs and ASICs clustering separately. However, it is not clear whether these clusters share functional characteristics. Indeed, the *C. elegans* DEG/ENaCs are the only group with members in multiple clusters. The vertebrate ASICs closely cluster with the TadNaCs from *Trichoplax adhaerens*, a primitive multicellular animal lacking any internal organs and neurons, and both groups have members that are sensitive to changes in proton concentration (Zhang & Canessa, 2002; Elkhatib *et al*., 2019).

**Figure 1:**
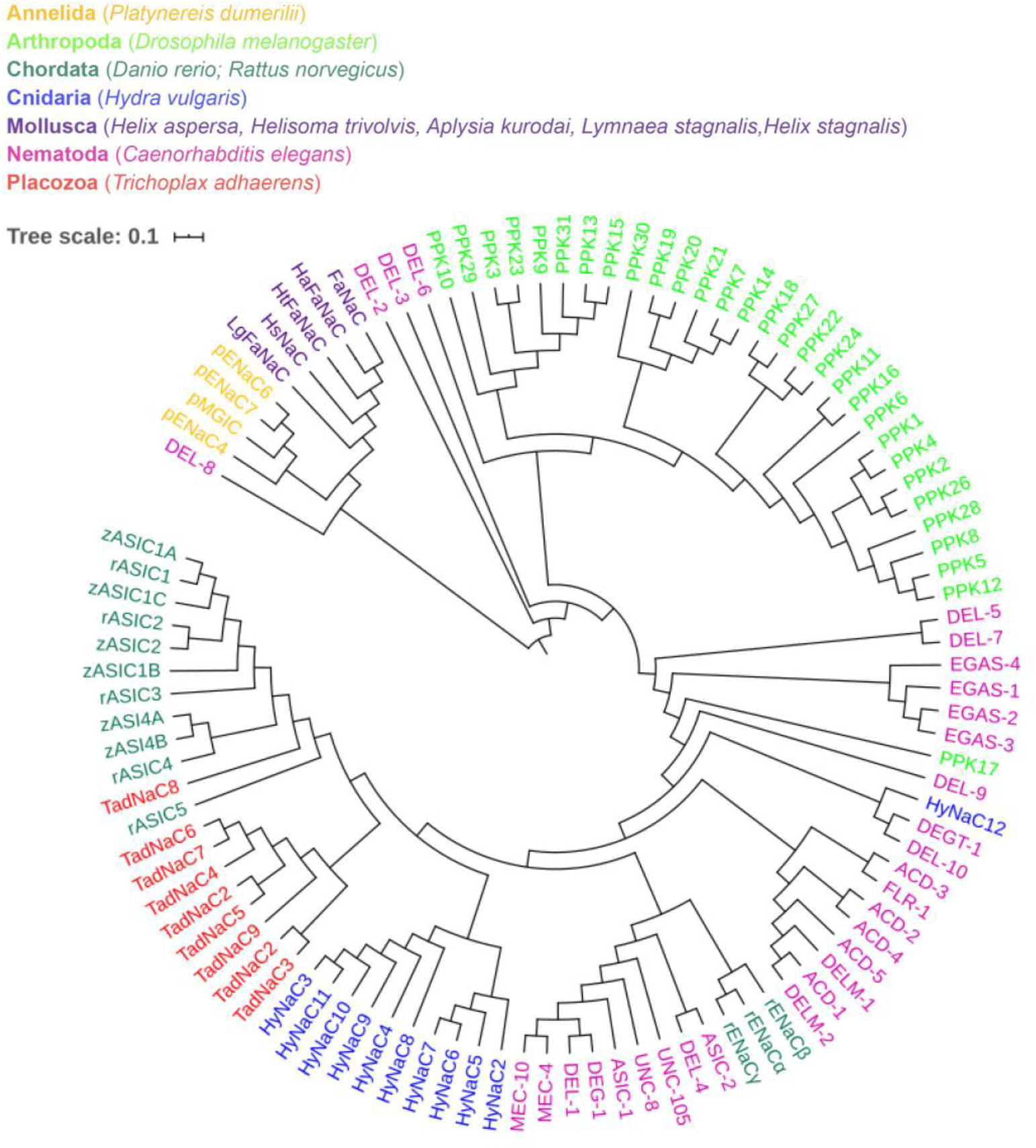
Phylogram of the DEG/ENaC family. Polar view of a phylogenetic tree of the DEG/ENaC super-family, aligned in MAFFT version 7 (see Methods for details). Tree scale represents the amount of genetic change. The data used to build these networks originate from the UniProt Consortium databases and the InterPro and ENA databases from EMBL-EBI. Colouring is according to phyla, as indicated. Accession numbers can be found in the Supplementary table 1.

**Figure 2:**
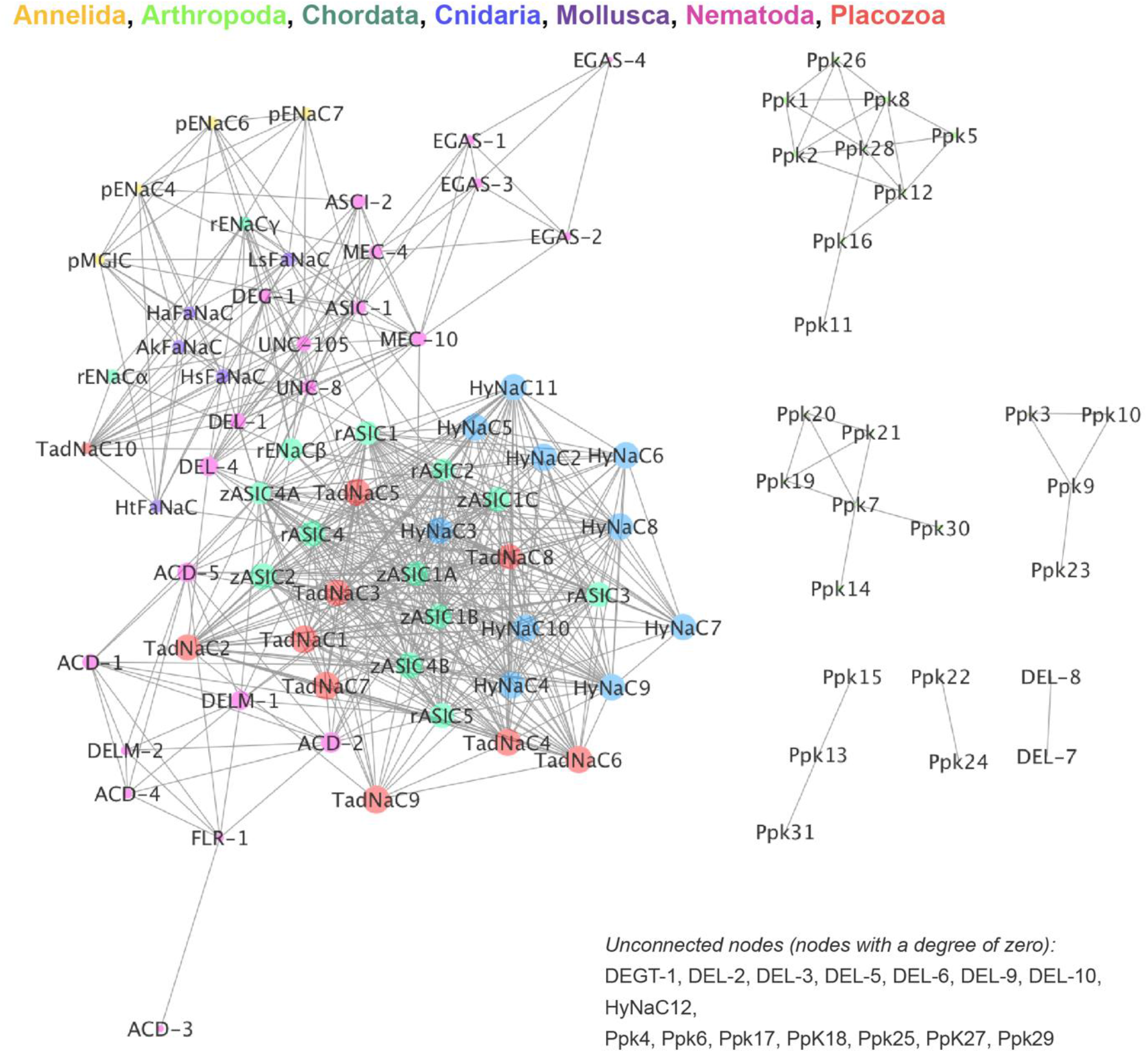
Sequence Similarity Network (SSN) of the DEG/ENaC family. Sequence Similarity Network (SSN) of diverse members of the DEG/ENaC family. Generated using the web tool for SSNs for protein families (EFI-EST) developed by the Enzyme Function Initiative efi.igb.illinois.edu/ (Gerlt *et al*., 2015; Gerlt, 2017; Zallot *et al*., 2018; Zallot *et al*., 2019). Each symbol represents a protein (node), two nodes are connected by a line (edge) if they share >25% sequence similarity and lengths of edges correlate with the relative dissimilarities of each pair. Relative positioning of disconnected clusters and nodes has no meaning. Unconnected notes, i.e. nodes with a degree of zero, are indicated on the bottom right-hand side. Cytoscape (Shannon *et al*., 2003) is used to explore SSNs. Node sizes are determined by the degree of connectivity of the nodes (number of edges). The EFI-EST webtools use NCBI BLAST and CD-HIT to create SSNs. The computationally-guided functional profiling tool uses the CGFP programs from the Balskus Lab (https://www.microbialchemist.com/metagenomic-profiling/) (Levin *et al*., 2017) and ShortBRED from the Huttenhower Lab (http://huttenhower.sph.harvard.edu/shortbred) (Kaminski *et al*., 2015). The data used to build these networks originate from the UniProt Consortium databases and the InterPro and ENA databases from EMBL-EBI. Node colouring is according to phyla, as indicated. Accession numbers can be found in the Supplementary table 1.

Despite the expansion and known diversity of *C. elegans* DEG/ENaCs, many members lack functional characterization at the level of the channel, an obvious limitation when interpreting their *in vivo* function. Therefore, we set out to perform a comprehensive screen for acid-sensitive *C. elegans* DEG/ENaC channel subunits. In addition to five subunits that form acid-inhibited homomers, we identified three acid-activated members. We demonstrate a diversity in the effect of amiloride, with some being blocked, but others being unaffected or potentiated, and also show that the acid-activated members exhibit diversity in ion selectivity. Like their mammalian counterparts, their currents are blocked or potentiated by zinc, indicating further conservation of function across phyla. This characterisation serves as a starting point for further screens for compounds that affect these channels and for understanding the molecular basis of diversity in DEG/ENaC function.

## Methods

### Protein sequences and alignment

Sequences of the DEG/ENaC superfamily were from UniProt, NBCI, Wormbase and Flybase, combined into one file using SnapGene (Insightful Science, snapgene.com). Due to the recent finding that removal of noisy or uncertain columns does not necessarily improve phylogenetic reconstruction (Tan *et al*., 2015), the complete amino acid sequences of the longest isoform (where applicable) were used for the phylogenetic estimation of protein similarity. To address the issue that variable regions tend to be over-aligned, and consequently, might lead to biases, the robust aligners PRANK ((Loytynoja & Goldman, 2010) data not shown) and MAFFT (Katoh *et al*., 2002; Katoh & Standley, 2013) were used and confidence in the individual alignment columns was assessed using GUIDANCE2. Both alignments generated were similar. We used MAFFT (Katoh *et al*., 2002; Katoh & Standley, 2013) as it allows re-adjustment alignment to reflect information from sequences aligned later (Larkin *et al*., 2007).

#### Phylogram

Visualisation of the DEG/ENaC super-family, protein sequences were aligned in MAFFT version 7 multiple alignment program using rough distance and average linkage UPGMA (unweighted pair group method with arithmetic mean) and the tree was visualized using iTOL (Ciccarelli *et al*., 2006; Kuraku *et al*., 2013; Letunic & Bork, 2019).

#### Sequence Similarity Network (SSN)

SSNs were generated using the web tool for SSNs for protein families (EFI-EST) developed by the Enzyme Function Initiative (EFI; efi.igb.illinois.edu/) (Gerlt *et al*., 2015; Gerlt, 2017; Zallot *et al*., 2018; Zallot *et al*., 2019). Cytoscape (Shannon *et al*., 2003) was used to explore SSNs. The EFI-EST webtools use NCBI BLAST and CD-HIT to generate SSNs. The computationally-guided functional profiling tool uses the CGFP programs from the Balskus Lab (https://bitbucket.org/biobakery/cgfp/src) (Levin *et al*., 2017) and ShortBRED from the Huttenhower Lab (http://huttenhower.sph.harvard.edu/shortbred) (Kaminski *et al*., 2015). The data used in these analyses originated from the UniProt Consortium databases and the InterPro and ENA databases from EMBL-EBI.

### Molecular Biology

The transcriptional reporters for the *Pdel-9::GFP* and *Pacd-2::GFP* plasmids were a kind gift from Professor Kyuhyung Kim’s lab (Daegu Gyeongbuk Institute of Science & Technology (DGIST), Korea) containing 3113 bp upstream of the *del-9* gene and 3004 bp *acd-2* promoter (2526 bp upstream of the start codon of *acd-2* and including 478 bp fragment of the *acd-2* gene). The *asic-1* promoter consisted of 3500 bp upstream of the start codon. All plasmids, including the fluorophore-reporter-fusions and cloning of cDNA into KSM vector for *Xenopus* oocyte expression, were assembled using NEBuilder^®^ HiFi DNA Assembly Master Mix (Catalogue # E2621L). *C. elegans* cDNA was obtained from growing N2 wild-type animals on fifteen 6cm NGM plates until the food was diminished, and subsequently extracted and purified using the TRIzol Direct-zol RNA Miniprep (Catalogue #R2051, Zymo Research). cDNA was generated using the Invitrogen™ SuperScript™ III First-Strand Synthesis System (Catalogue # 18080051). Primers were designed using SnapGene 5.0.4. (Hifi-Cloning of two fragments) based on the cDNA gene sequence found on wormbase.org, and ordered from Integrated DNA Technologies Inc (IDT) (Leuven Belgium) or Sigma-Aldrich (Merck Life Science UK Limited, Dorset, UK). The cDNA inserts were sub-cloned into the KSM vector under the control of the T7 promoter, with 5’ and 3’ untranslated regions (UTRs) of the *Xenopus* beta-globin gene and a poly(A) tail. The forward primer *agatctggttaccactaaaccagcc* and reverse primer *tgcaggaattcgatatcaagcttatcgatacc* were used to amplify the KSM vector. NEB Tm Calculator was used to determine annealing temperatures.

### Two-Electrode Voltage Clamp (TEVC) in Xenopus oocytes

cRNA was synthesized using the mMessage mMachine T3 Transcription Kit (Ambion # AM1348), purified with GeneJET RNA Cleanup and Concentration Micro Kit (Thermo Scientific # K0841) and eluted in 15μL RNase free water. *Xenopus laevis* oocytes of at least 1mm in size were obtained from EcoCyte Bioscience (Dortmund, Germany). They were de-folliculated by collagenase treatment and maintained in standard 1XND96 solution (96mM NaCl, 2mM MgCl2, 5mM HEPES, 2mM KCl, 1.8mM CaCl2, pH7.4). Oocytes were injected with 25μL of cRNA solution at a total concentration of approximately 500 ng/μL using the Roboinject (MultiChannel Systems). Oocytes were kept at 16°C in 1XND96 prior to TEVC. TEVC was performed 1-2 days post-injection at room temperature using the Roboocyte2 (MultiChannel Systems). *Xenopus* oocytes were clamped at −60 mV, using ready-to-use Measuring Heads from MultiChannel Systems filled with 1.0 M KCl and 1.5 M K-acetate. All channels were tested using the Roboocyte2 (MultiChannel Systems). For all current-voltage steps (I-V) experiments, measurements were obtained in each solution once a steady-state current was achieved and the background leak current was subtracted.

As millimolar concentrations of Ca^2+^ and other divalent ions except Mg^2+^ can block ASIC currents (Paukert *et al*., 2004), Ca^2+^-free buffers were used for substitution experiments of monovalent cations, adapted from a previous protocol (Hardege *et al*., 2015): 96mM XCl, 1mM MgCl2, 5mM HEPES, pH adjusted to 7.4 with XOH, where X was Na^+^, K^+^ or Li^+^, respectively. The osmolarity was checked and confirmed to be within the error of 210 mosm or adjusted with D-Glucose if necessary (Awayda & Subramanyam, 1998). For testing pH sensitivity, 1x ND96 solutions was used; for solutions with a pH 5 or lower, MES was used instead of HEPES and adjusted with HCl. Actual current I-V curves for each individual oocyte were fitted to a linear regression line and the x intercept was compared between solutions to calculate an average reversal potential (Erev). Reversal potential shift (ΔErev) when shifting from a NaCl to a KCl or LiCl solution was calculated for each individual oocyte. In order to test the responses to pH, the channel-expressing *Xenopus* oocytes were perfused with 1XND96 (using HEPES for buffering pH above 5.5 and MES for pH below 5), pH was adjusted with HCl and ranged from pH 7.4 (neutral pH of the ND96 solution) to pH 4. For the Zn^2+^ dose-responses, a 1 M ZnCl2 stock solution in water (Catalogue #229997, Sigma-Aldrich) was diluted to the desired concentrations in ND96 buffer based on a previously established protocol (Chen *et al*., 2012). Zn^2+^ was applied at increasing concentrations in the range of 0.1 μM to 5 mM. Baseline subtraction and drift correction for all dose-responses was applied in the Roboocyte2+ software (MultiChannel Systems). For analysis, currents were normalized to maximal currents (I/I_max_) and best fitted with the Hill equation (Variable slope) in GraphPad Prism version 9.0.2.

### Confocal microscopy

Worms were mounted on 3% agar pads (in M9 (3 g KH2PO4, 6 g Na2HPO4, 5 g NaCl, 1M MgSO4)) in a 3 μL drop of M9 containing 25 mM sodium azide (NaN3, Sigma-Aldrich). Images were acquired using a Leica TCS SP8 STED 3X confocal microscope at 63x, 40x or 20x resolution and Z stacks were generated using Fiji (ImageJ) (Schneider et al., 2012).

### Availability of materials

*C. elegans* strains and plasmids generated for the purpose of this study are available upon request from.

## Results

### Identification of two groups of acid-sensitive C. elegans DEG/ENaCs

In contrast to the mammalian ASICs, which are activated in response to a drop in pH, ACD-1, the only *C. elegans* DEG/ENaC for which proton sensitivity had been characterised in *Xenopus* oocytes, is open at neutral pH and inhibited by low pH (Wang *et al*., 2008). To identify other acid-sensing *C. elegans* DEG/ENaC members, we performed a systematic screen for pH-sensitive subunits using Two-Electrode Voltage Clamp (TEVC) in *Xenopus* oocytes injected with cRNA, for all 30 *C. elegans* DEG/ENaCs. For the initial screen, oocytes were perfused with pH 7.4 and pH 4 solutions, and currents were recorded and quantified (Figure 3A-E). We did not go below pH 4 for the screen as our control *Xenopus* oocytes injected with nuclease free water started responding to solutions below pH 4. We identified two groups of acid-sensing DEG/ENaCs. One group included the previously characterised ACD-1 (Wang *et al*., 2008) (Figure 3B), showing currents at neutral pH (pH 7.4) that were decreased at low pH (pH 4). We identified four additional members, ACD-5, DEL-4 DELM-1 and UNC-105 (isoform h), which also fit this profile and which we will refer to here as acid-inhibited channels (Figure 3D). Interestingly, we also found three members that showed robustly increased currents at low pH, i.e. that were opened, or further opened, in response to increasing proton concentrations; these were ASIC-1, ACD-2 and DEL-9 (Figure 3C). We designated the members in this second group as acid-activated channels.

**Figure 3:**
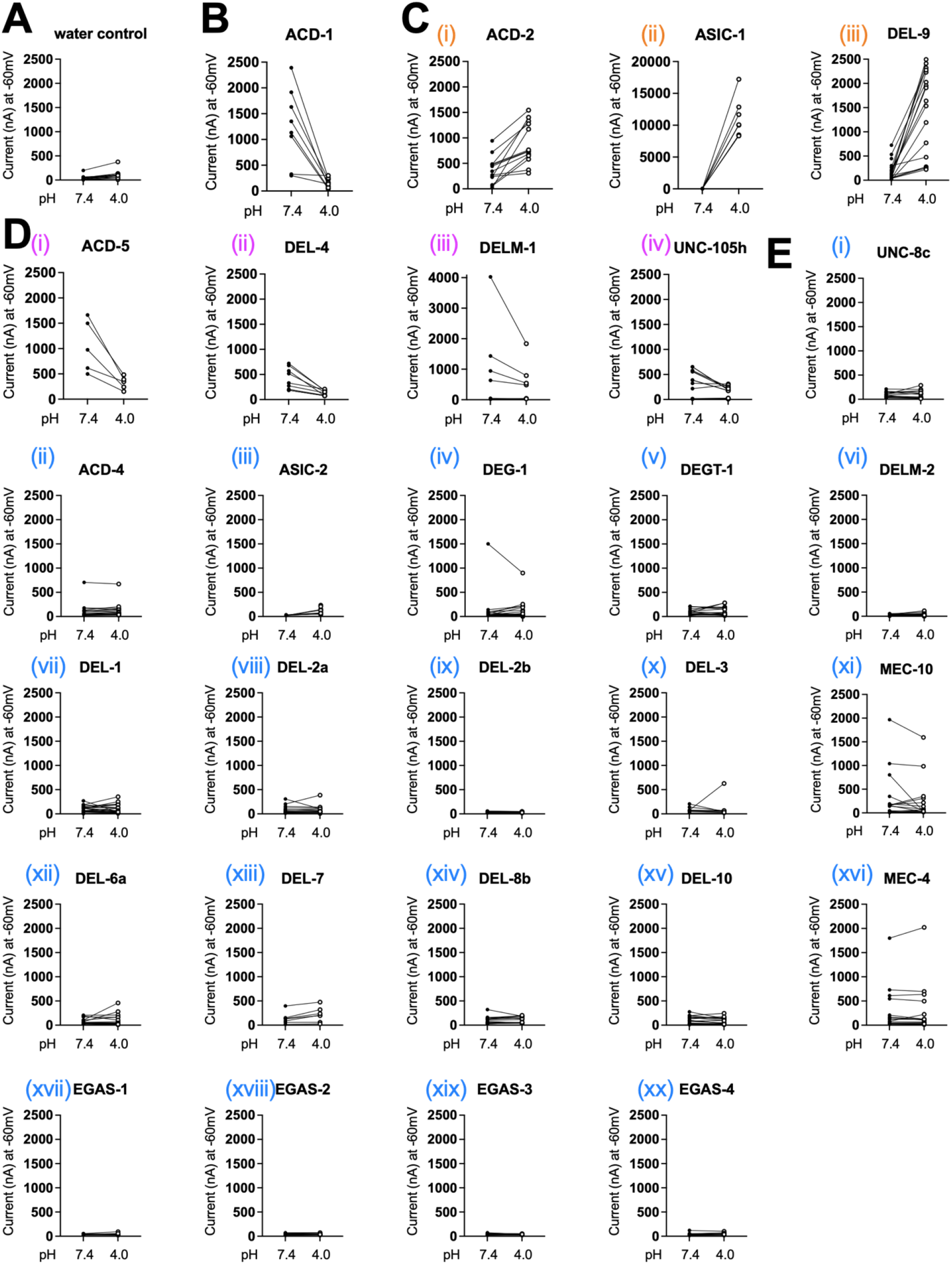
*Quantification of current at pH 7.4 and pH 4 of Xenopus oocytes* expressing *C. elegans DEG/ENaC subunits*. *Xenopus* oocytes injected with the respective cRNA. Lines connect data from individual oocytes. (A) Nuclease-free water injected oocytes serve as negative controls (n=13). (B) ACD-1 (n=8) expressing oocytes serve as a positive control (Wang *et al*., 2008). (C) Constructs that form acid-activated channels: ACD-2 (n=13), ASIC-1 (n=7) and DEL-9 (n=19). (D) Constructs that form acid-inhibited channels: ACD-5 (n=5), DEL-4 (n=9), DELM-1 (n=7), UNC-105 (n=17). (E) *Xenopus* oocytes injected with the respective cRNA do not display acid-sensitive currents: UNC-8c (n=14), ACD-4 (n=23), ASIC-2 (n=21), DEG-1 (n=23), DEGT-1 (n=23), DELM-2 (n=24), DEL-1 (n=24), DEL-2a (n=18), DEL-2b (n=10), DEL-3 (n=10), MEC-10 (n=17), DEL-6a (n=23), DEL-7 (n=7), DEL-8b (n=20), DEL-10 (n=18), MEC-4 (n=16), EGAS-1 (n=12), EGAS-2 (n=12), EGAS-3 (n=11), EGAS-4 (n=12). The pH-insensitive currents of ACD-3 and DEL-5 are published elsewhere (Kaulich *et al*., 2021). Axis labels: y= actual current in nA; x = perfusion with either pH 7.4 (closed circle) or pH 4 (open circle) as indicated. Holding potential was - 60mV. Experiments were repeated more than three times and on different days and were pooled together. Note that all graphs are plotted using the same scale, except ASIC-1 and DELM-1, due to the large currents observed in oocytes expressing these two constructs.

### The C. elegans DEG/ENaCs include both acid-inhibited and acid-activated channels

To characterise the acid-sensitivity and ion selectivity of the acid-sensitive DEG/ENaCs identified in the screen, we perfused different pH solutions over the oocytes expressing the respective cRNA and substituted different monovalent cations in the recording solution. The ion selectivity experiments were performed at pH_50_ concentrations and the shift in reversal potential (ΔErev) was assessed after substituting NaCl with either KCl or LiCl in the solution containing 96mM NaCl, 1mM MgCl2, 5mM MES, based on a previous protocol (Hardege et al., 2015). The osmolarity was adjusted to 210 mosm using D-Glucose (Awayda & Subramanyam, 1998). All ion-substitution experiments were conducted in the absence of Ca^2+^ as Ca^2+^ has been shown to be able to block ASIC channels (Paukert et al., 2004).

We first investigated the properties of the acid-inhibited channels. ACD-5, which we have described elsewhere, was permeable to Li^+^ and to a lesser degree to Na^+^ and K^+^ but impermeable for Ca^2+^ and exhibited maximal current at approximately pH 6 and was inhibited by both lower and higher pH, with pH_50_s of 4.87 and 6.48 (Kaulich *et al*., 2021). In contrast, DEL-4 was shown to be a Na^+^ channel that can be blocked in the range of pH 8-4 with pH_50_ of 5.7 (Petratou *et al*., submitted). DELM-1 had previously been characterised as an amiloride-sensitive channel permeable for Li^+^ and Na^+^, and to a lesser degree to K^+^ (Han *et al*., 2013). We found that DELM-1 was inhibited by acidic pH with a pH_50_ of 5.75 (Figure 4A, B). The Na^+^ and K^+^ permeable UNC-105 has previously been implicated in proton-sensing *in vivo* in muscles (Jospin & Allard, 2004). We found that UNC-105 (isoform h) produced small currents which were inhibited when perfused with pH in the range of pH 9-4 with a pH_50_ of 6.42 (Figure 4A, B).

**Figure 4:**
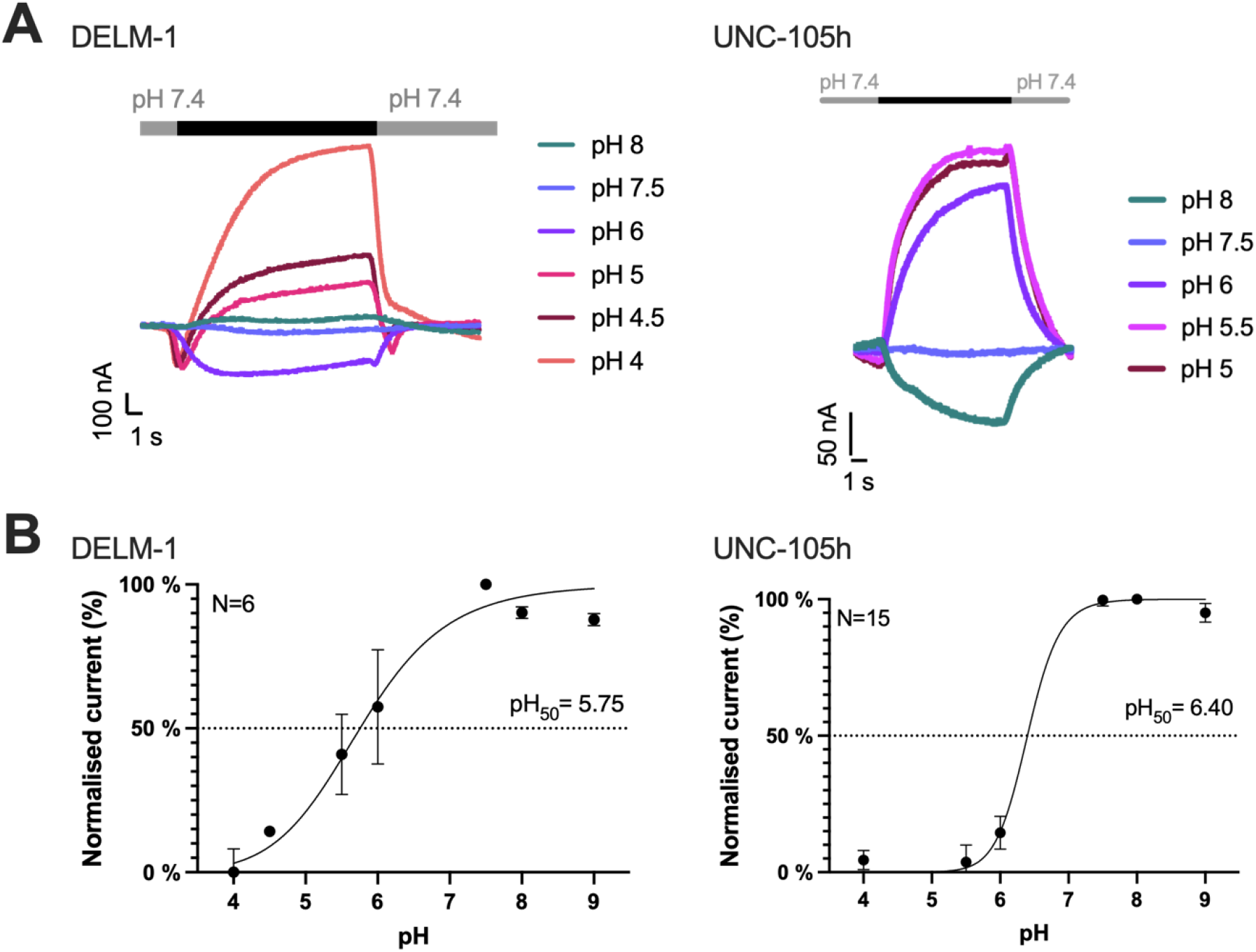
DELM-1 and UNC-105 form acid-inhibited homomeric channels. (A) Representative traces of *Xenopus* oocytes expressing the respective construct when perfused with solutions of different pH, lowering of pH (black bar) from a holding pH of 7.4. (B) Current-pH relationship. Dotted line indicates pH_50_. DELM-1, pH_50_ = 5.75 ± 0.23 (N = 6). UNC-105 isoform h, pH_50_ = 6.42 ± 0.10 (N = 15). Currents were recorded at a holding potential of – 60mV, normalized to maximal currents (I/I_max_) and best fitted with the Hill equation (Variable slope). Error bars represent Mean ± SEM.

By contrast, for all three acid-activated subunits, acid-evoked currents increased in a concentration-dependent manner. We observed an excitatory pH_50_ of 4.49 for ASIC-1, pH_50_ of 5.03 for ACD-2 and pH_50_ of 4.27 for DEL-9. (Figure 5A, B). None of the three homomeric channels desensitized, but reached a plateau after several seconds, with ASIC-1 and ACD-2 reaching the plateau much faster than DEL-9, as shown by the timescale in Figure 5A, B. This is not an artefact of the perfusion rate as the same perfusion rate was used for all oocytes and conditions. ACD-2 partially desensitised at pH 4 (and below, Supplementary figure S1), which might be relevant to the channel’s physiological pH range.

**Figure 5:**
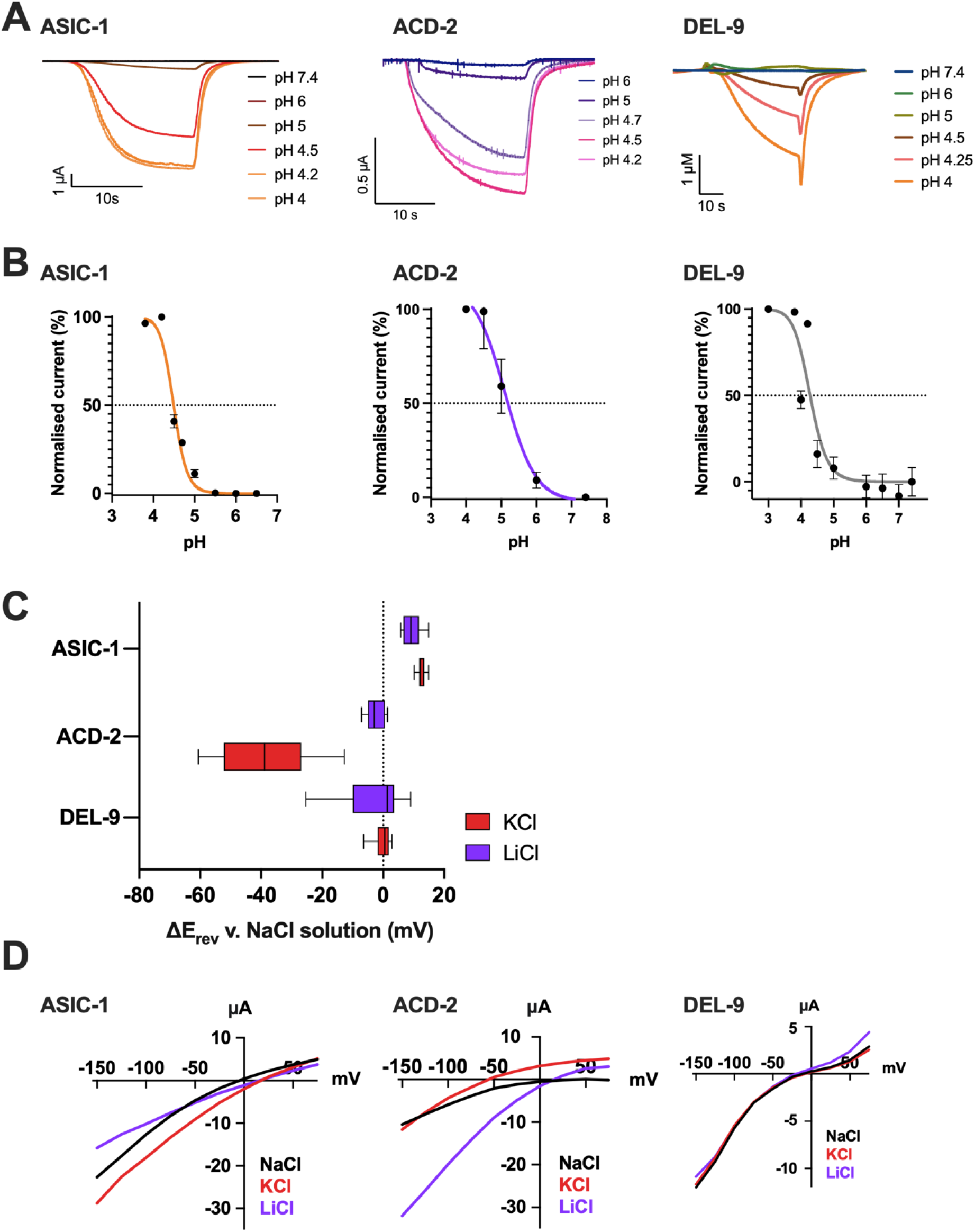
ASIC-1, ACD-2 and DEL-9 form acid-activated homomeric channels permeable for monomeric cations. (A) Representative traces of *Xenopus* oocytes expressing the respective construct when perfused with solutions of different pH, lowering of pH from a holding pH of 7.4. (B) Current-pH relationship. Dotted line indicates pH_50_. ASIC-1, pH_50_ = 4.49 ± 0.02 (N = 13). ACD-2, pH_50_ = 5.028 ± 0.109 (N = 6). DEL-9, pH_50_ = 4.27 ± 0.07 (N = 10). Currents were recorded at a holding potential of – 60mV, normalized to maximal currents (I/I_max_) and best fitted with the Hill equation (Variable slope). Error bars represent Mean ± SEM. (C-D) Summary of Ion selectivity. ASIC-1, ACD-2 and DEL-9 are permeable for monomeric cations. (C) Average calculated from 4<N<12 oocytes for each construct of ΔErev when shifting from a NaCl solution to KCl or LiCl solution. A negative shift of E_rev_ indicating a preference for Na^+^ over the respective ion and a positive shift indicating a preference of the respective ion over Na^+^. Data is presented as box-plots (hinges of the plot are the 25th to 75th percentiles) with Median and whiskers of 1.5IQD (inter-quartile distance, as calculated by the Tukey method). (D) Representative current-voltage (IV) relationships for oocytes expressing ASIC-1, ACD-2 and DEL-9. Actual current for each oocyte and baseline current subtracted from acid-evoked current at pH_50_ concentrations (see panel B for reference). The oocyte membrane was clamped at −60 mV and voltage steps from −150 mV to +75 mV were applied as indicated. Currents are actual currents in μA (y-axis), voltage steps in mV (x-axis) as indicated. Error bars represent Mean ± SEM.

As no physiological data was available for any of the three acid-activated DEG/ENaC subunits, we investigated their ion selectivity by carrying out ion substitution experiments. Our results showed that ASIC-1 exhibits a preference of K^+^ and Li^*+*^ over Na^+^, with a median positive shift in E_rev_ of 12.2 mV, when shifting from a NaCl solution to a KCl or LiCl solution (Figure 5C, D). In contrast ACD-2 was observed to be a sodium channel, selective for Li^+^ and Na^+^ over K^+^ with no change in E_rev_ when switching from a NaCl to a LiCl solution, but a large median negative shift in E_rev_ of −43.3 mV when shifting from a NaCl solution to a KCl solution (Figure 5C, D). This ion selectivity property is similar to that previously described for DELM-1 (Han *et al*., 2013). Finally, DEL-9 was non-selective for monovalent cations, showing no shift in reversal potential when switching between solutions (Figure 5C, D). Thus, the newly characterised acid-sensitive DEG/ENaCs showed significant diversity in their ion selectivity properties, consistent with previous findings on other family members (Canessa *et al*., 1994; Wang *et al*., 2008; Carattino & Della Vecchia, 2012; Fechner *et al*., 2021).

### Exploring the effect of known DEG/ENaC modulators amiloride and zinc

Both amiloride and zinc can act as blockers for some DEG/EnaCs and as enhancers for others, again showing the functional diversity of this channel family. Amiloride is an anti-hypertensive drug, acting on the human ENaCs in the kidney (Teiwes & Toto, 2007; Bhagatwala *et al*., 2014). It is a potent blocker of many, but not all DEG/ENaCs. Previous work showed that homomeric ASIC3, ASIC2 and heteromeric ASIC3/ASIC1b are potentiated by amiloride, whereas *C. elegans* DEGT-1 is amiloride-insensitive (Bentley, 1968; Palmer & Frindt, 1986; Canessa *et al*., 1994; Schild *et al*., 1997; Adams *et al*., 1999; Kellenberger *et al*., 2003; Li *et al*., 2011; Baconguis *et al*., 2014; Besson *et al*., 2017; Fechner *et al*., 2021; Matasic *et al*., 2021). Therefore, we investigated if the acid-evoked currents at the respective pH_50_ of ASIC-1, ACD-2 and DEL-9 could be blocked by or potentiated by amiloride (Figure 6; for pH_50_ see Figure 5B for reference). We observed that ASIC-1 and ACD-2 are sensitive to amiloride; indeed acid-evoked currents could be blocked in a dose-dependent manner, which is also a common characteristic of the DEG/ENaC super-family (Vullo & Kellenberger, 2020). Amiloride blocked ASIC-1 acid-evoked currents with an IC_50_ of 134 μM (Figure 6B) and ACD-2 acid-evoked currents with an IC_50_ of 84 μM (Figure 6C). By contrast, DEL-9 acid-activated currents were insensitive to amiloride (Figure 6D).

**Figure 6:**
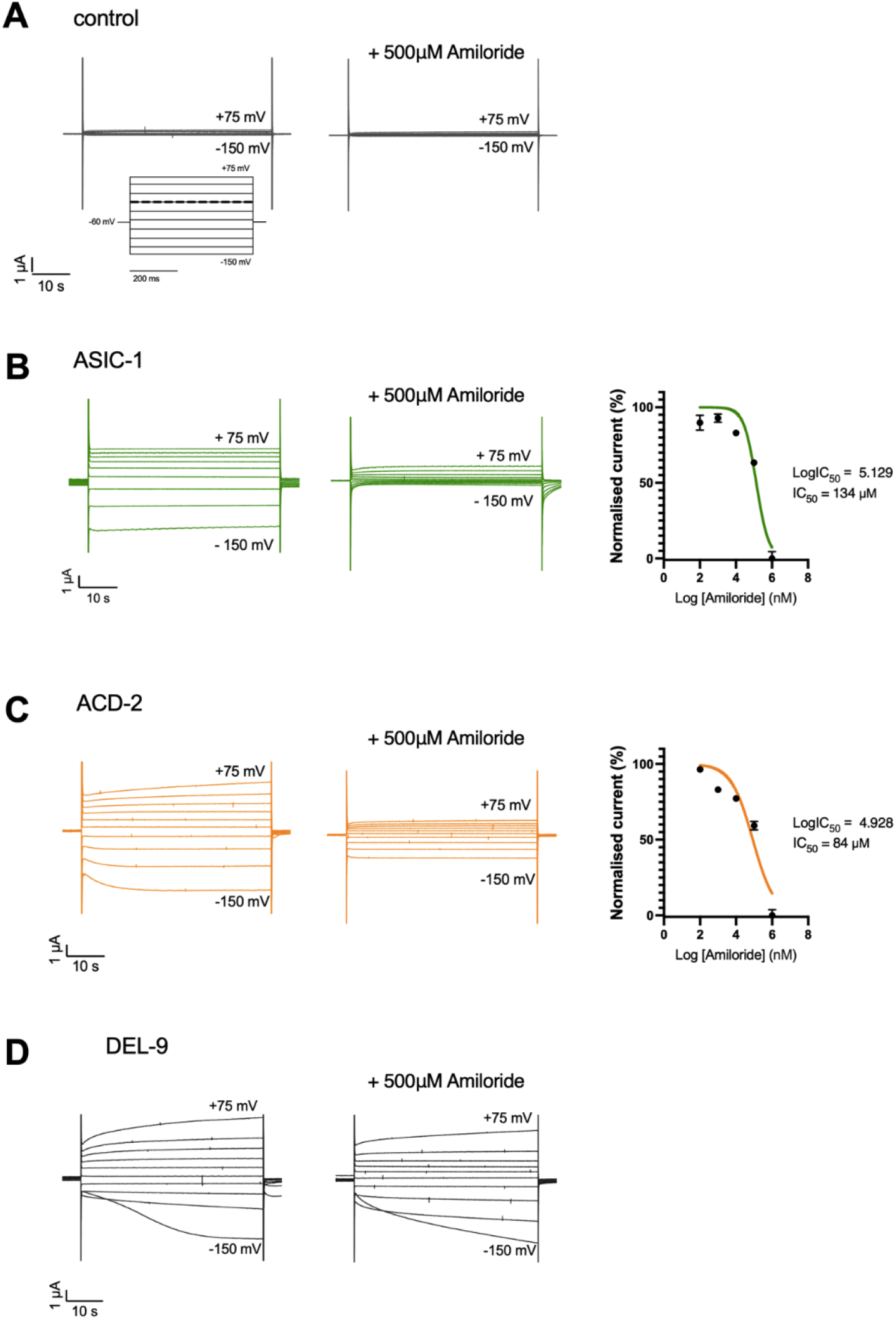
Amiloride-sensitivity ACD-2, ASIC-1 and DEL-9. ASIC-1 and ACD-2 (but not DEL-9) acid-evoked transient currents can be blocked by amiloride. Representative acid-evoked transient currents in the absence (left) and presence of 500 μM amiloride (middle) at pH_50_ (for pH_50_ of each channel see Figure 5B). (A) Nuclease-free water-injected oocytes (negative control) are unaffected by low pH (example shown here pH 4.5) or amiloride. (B) Amiloride dose-response of ASIC-1 with an IC_50_ of 134 μM amiloride (N = 10) and (C) ACD-2 with an IC_50_ of 84 μM amiloride (N = 12). (D) DEL-9 (N=10) acid-activated transient currents are unaffected by amiloride. Currents were recorded at a holding potential of - 60mV and are normalised to the maximum current (I_max_) calculated for each oocyte individually, and best fitted with the Hill equation (Variable slope). The *Xenopus* oocyte membrane was clamped at −60 mV and voltage steps from −150 mV to +75 mV were applied as indicated. Currents are actual currents in μA (y-axis), voltage steps in mV (x-axis) as indicated. Error bars represent Mean ± SEM.

We also investigated the effect of amiloride on the acid-inhibited subunits, ACD-1, DELM-1 and UNC-105, ACD-5 and DEL-4. We observed that all were inhibited by amiloride (Supplementary figure 2, see also (Jospin & Allard, 2004; Wang *et al*., 2008; Han *et al*., 2013; Kaulich *et al*., 2021) (Petratou *et al*., submitted)). UNC-105 showed paradoxical block for amiloride with an EC_50_ of 14.56 μM amiloride and a strong inhibition at 1 μM (Figure 7 A, B).

**Figure 7:**
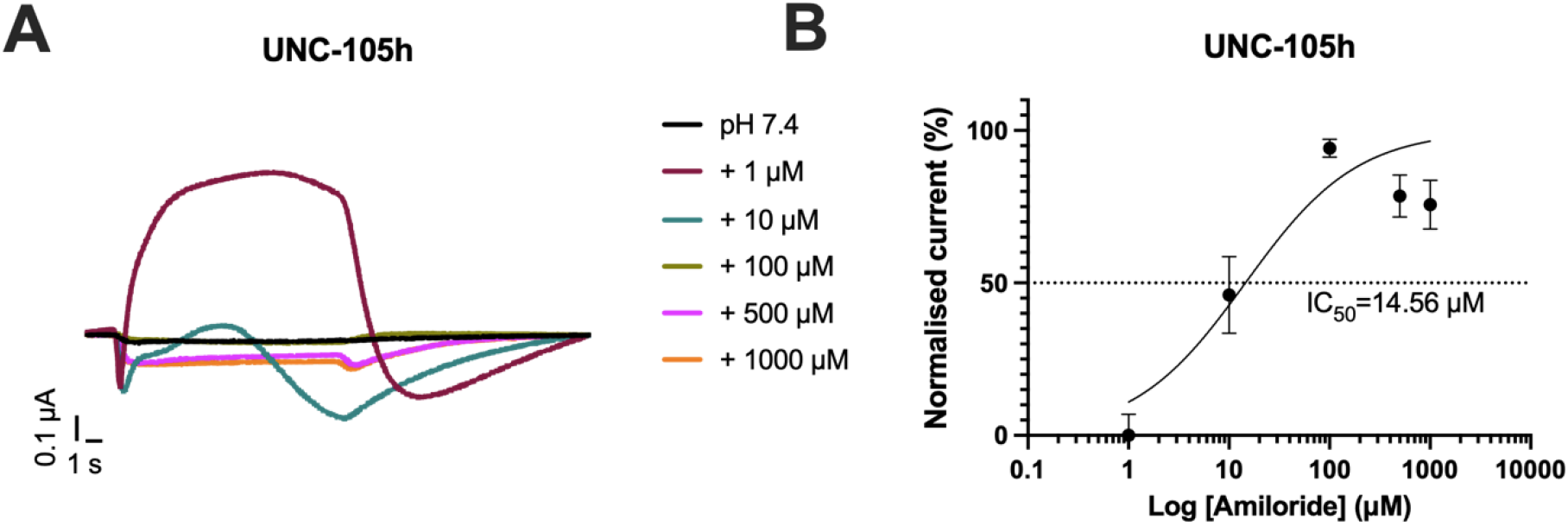
Paradoxical effect of amiloride on UNC-105. UNC-105 (isoform h) transient currents at pH 7.4 can be blocked by amiloride. (A) Representative transient currents at increasing concentrations of amiloride. Amiloride blocks the channel at 1 μM concentration but other concentrations do not affect the currents. The *Xenopus* oocyte membrane was clamped at −60 mV. Currents are actual currents in μA (y-axis), voltage steps in mV (x-axis) as indicated. (B) Amiloride dose-response of UNC-105 with an EC_50_ of 14.56 μM amiloride (N = 16). Currents were recorded at a holding potential of - 60mV and are normalised to the maximum current (I_max_) calculated for each oocyte individually, and best fitted with the Hill equation (Variable slope). Error bars represent Mean ± SEM.

From the screen shown in Figure 3, we found other DEG/ENaC subunits exhibiting a current at pH 7.4. This “leak” current at pH 7.4 could be an indicator of an open channel, which may or may not be constitutively open in its physiological context, depending on the pH range of its environment, or it might be independent of pH-sensitivity. However, a leak current could also potentially result from non-functional channel expression making the oocytes unhealthy. Since amiloride can have either a potentiating or inhibitory effect on the channel’s current, we also examined amiloride sensitivity for all other DEG/ENaC channels. We found that whereas most channels’ currents were unaffected by amiloride and the currents at pH 7.4 persisted (Supplementary figures 3 and 4), DEL-5, DEL-8b and DEL-9 currents at pH 7.4 were significantly potentiated by amiloride (see also (Kaulich *et al*., 2021)) (Supplementary figure 5). Thus, the effect of amiloride on channel activity varied widely between different members of the *C. elegans* DEG/ENaC family.

Zinc is an essential trace element in the brain that is present in synaptic vesicles in a subgroup of glutamatergic neurons and co-released with glutamate in response to high-frequency stimulation (Assaf & Chung, 1984; Howell *et al*., 1984; Frederickson & Moncrieff, 1994). High concentrations of Zn^2+^ potentiate acid-evoked currents of homomeric and heteromeric ASIC2a-containing channels (Baron *et al*., 2001). By contrast, Zn^2+^ can block ASIC1b subunits, in which multiple binding sites for Zn^2+^ in the extracellular domain have been proposed (Baron *et al*., 2001; Jiang *et al*., 2012). We therefore investigated the effect of Zn^2+^ on the newly characterised acid-sensing channels. We found that the pH_50_-evoked transient currents of the acid-activated channels could be blocked by Zn^2+^ in a dose-dependent manner, with an IC_50_ of 132.4 μM Zn^2+^ for ASIC-1, an IC_50_ of 46.39 μM Zn^2+^ for ACD-2, and an IC_50_ of 58.33 μM Zn^2+^ for DEL-9 (Figure 8). We further tested Zn^2+^ modulation at pH 7.4 (which corresponds to the open state of the channels) of the acid-inhibited channels and found that similar to the acid-activated channels, the acid-inhibited ACD-1, ACD-5, DEL-4 and UNC-105 were blocked by increasing concentrations of Zn^2+^ with IC_50_s of 184.4 μM, 165 μM, 12.25 μM and 31.94 μM, respectively (Figure 9A, B, D, E). In contrast, increasing concentrations of Zn^2+^ potentiated the baseline currents of DELM-1 with an EC_50_ of 238.3 μM (Figure 9C). This demonstrates that like their mammalian counterparts, those acid-sensing DEG/ENaCs can be modulated by Zn^2+^ in an inhibiting or potentiating manner. This demonstrates that *C. elegans* acid-sensing DEG/ENaCs share the function of zinc- and amiloride-modulation with their vertebrate homologues.

**Figure 8:**
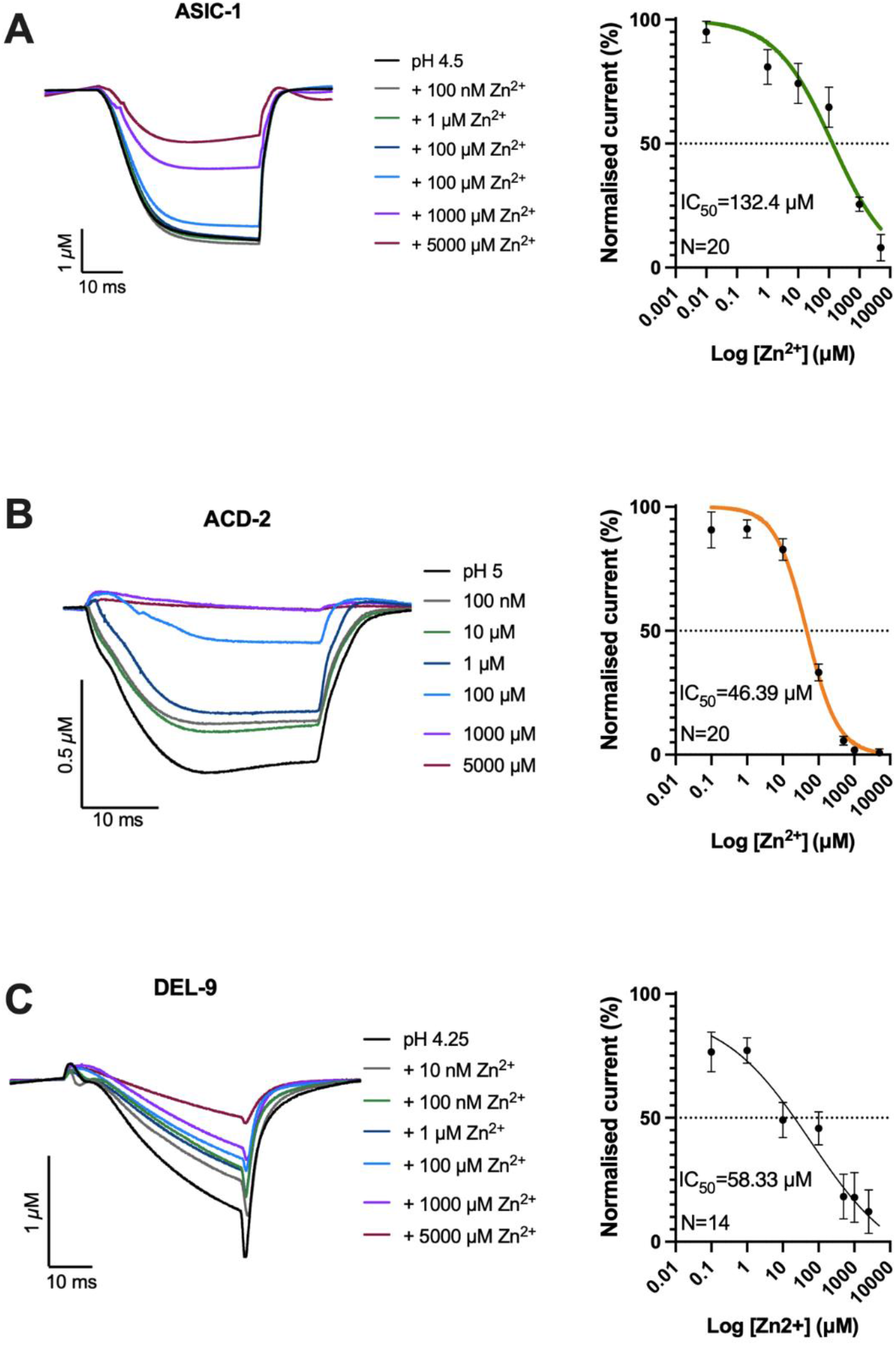
Zinc modulation of acid-activated DEG/ENaC channels in Xenopus oocytes. Zn^2+^ can block homomeric ASIC-1, ACD-2 and DEL-9 acid-evoked currents. Shown are representative example traces (Left) and dose response curves (Right) for each channel. (A) ASCI-1 (N=20), (B) ACD-2 (N=20) and (C) DEL-9 (N=14) pH_50_-evoked transient currents can be blocked by Zn^2+^ in a dose-dependent manner (at pH_50_ of each channel). Zn^2+^ dose-response of ASIC-1 with an IC_50_ of 132.4 μM Zn^2+^, ACD-2 with an IC_50_ of 46.39 μM Zn^2+^ and DEL-9 with an IC_50_ of 58.33 μM Zn^2+^. Baseline subtraction and drift correction was applied in the Roobocye2+ software. Currents were recorded at a holding potential of - 60mV and are normalised to the maximum current (I_max_) calculated for each oocyte individually, and best fitted with the Hill equation (Variable slope). Error bars represent Mean ± SEM.

**Figure 9:**
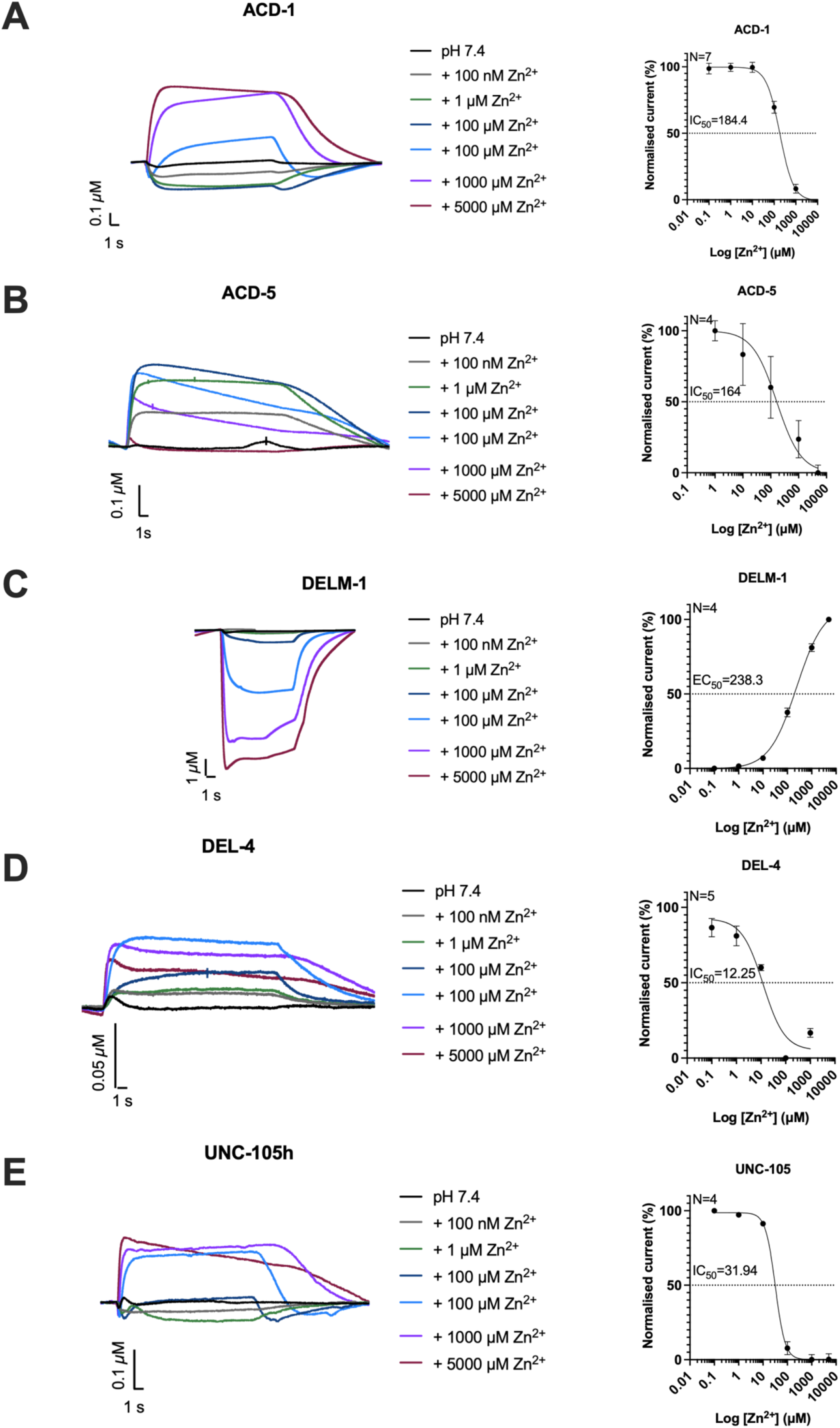
Zinc modulation of acid-inhibited DEG/ENaC channels in Xenopus oocytes. (A-E) Zn^2+^ modulate homomeric ACD-1, ACD-5, DELM-1, DEL −4 and UNC-105 currents at pH 7.4 in a dose-dependent manner. Shown are representative example traces (Left) and dose response curves (Right) for each channel: Zn^2+^ dose-response of ACD-1 with an IC_50_ of 184.4 μM Zn^2+^, ACD-5 with an IC_50_ of 165 μM, DELM-1 with an EC_50_ of 238.3 μM, DEL-4 with an IC_50_ of 12.25 μM, and DEL-4 with an IC_50_ of 31.94 μM Zn^2+^. Baseline subtraction and drift correction was applied in the Roobocye2+ software. Currents were recorded at a holding potential of - 60mV and are normalised to the maximum current (I_max_) calculated for each oocyte individually, and best fitted with the Hill equation (Variable slope). Error bars represent Mean ± SEM.

### Acid-activated DEG/ENaCs are expressed in both neurons and muscles

*C. elegans* DEG/ENaC expression has been described in various tissues including muscle, neurons, glia, and intestinal epithelia (Park & Horvitz, 1986a; Chalfie & Wolinsky, 1990; Driscoll & Chalfie, 1991; Huang *et al*., 1995; Take-Uchi *et al*., 1998; Wang *et al*., 2008; Wang *et al*., 2012; Han *et al*., 2013). The previously-described acid-inhibited DEG/ENaCs are expressed mainly in non-neuronal tissue with the exception of DEL-4 which is expressed in neurons (Petratou *et al*., submitted); ACD-1 and DELM-1 and DELM-2 are expressed in glia, ACD-5 and FLR-1 in the intestine, and UNC-105 is expressed in the body-wall muscle (Park & Horvitz, 1986a; Take-Uchi *et al*., 1998; Wang *et al*., 2008; Han *et al*., 2013; Kaulich *et al*., 2021).

To characterise the expression patterns of the acid-activated *C. elegans* DEG/ENaCs, we used transcriptional reporters. We confirmed previous reports that the *asic-1* promoter drives expression in the ADE, CEP, PVQ, PDE and PVD neurons (Voglis & Tavernarakis, 2008; Husson *et al*., 2012; De Stasio *et al*., 2018) and also observed expression in FLP and ventral cord neurons (Figure 10). The *acd-2* transcriptional reporter showed low-level expression in unidentified anterior neurons or glia in the head. Our *del-9* reporter was expressed in anterior and posterior body wall muscles, egg-laying muscles as well as head and tail neurons including AVL and PVQ. These and previous results show that the *C. elegans* acid-sensing DEG/ENaCs are not confined to a particular tissue but can be expressed in both neuronal and non-neuronal tissues.

**Figure 10:**
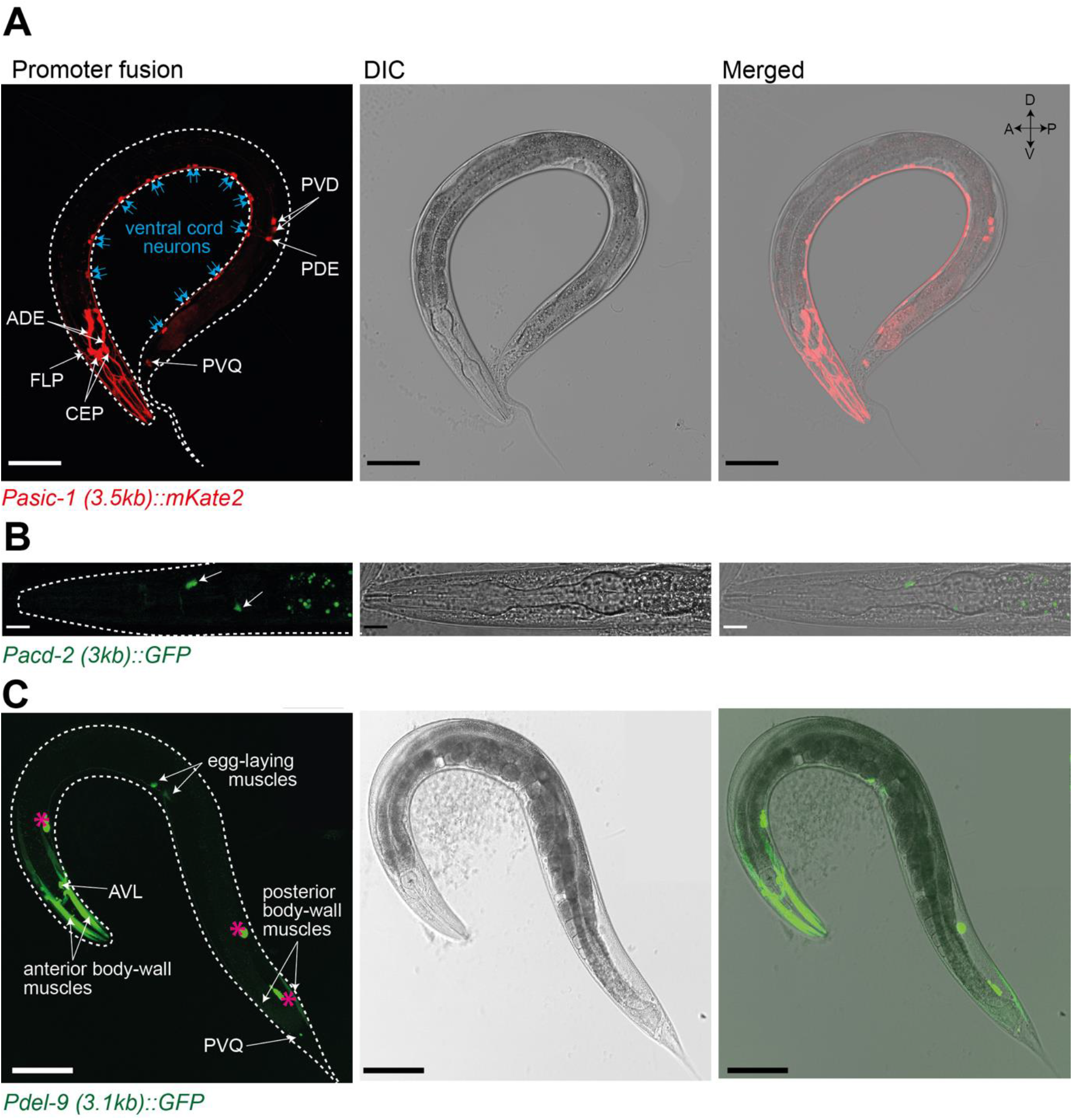
Acid-activated DEG/ENaCs are expressed in neuronal and non-neuronal tissue. Expression pattern of the transcriptional promoter fusions of the acid-activated DEG/ENaC genes with *mKate2* or *GFP* in L4 and young adults. (A) The *asic-1* promoter drives expression in the dopaminergic neurons and PVDs and FLPs. (B) *acd-2* promoter expression can be faintly detected in the head which based on localisation and RNA-sequencing data by (Cao *et al*., 2017) could be neurons or glia cells. Green dots in the intestine are autofluorescence. (C) The *del-9* promoter drives expression in the body-wall and egg-laying muscles, in head neurons, PVQ neuron and the GABAergic neuron AVL. Pink asterisks indicate the coelomocytes (*Punc-122::GFP*, used as co-injection marker to select transgenic animals). Scale bars: (A) 100 μm (B) 10 μm (C) 100 μm.

**Table 1:** Summary of physiological properties of C. elegans acid-sensing DEG/ENaCs.

|  |  | pH<sub>50</sub> | Ion selectivity | Amiloride | Zn<sup>2+</sup> | Expression | Reference |
| --- | --- | --- | --- | --- | --- | --- | --- |
| acid- inhibited | ACD-1 | 6.4 | Na<sup>+</sup>>Li<sup>+</sup>>K<sup>+</sup> | IC<sub>50</sub>=99<br>μM | EC<sub>50</sub>=184.4<br>μM | glia | (Wang *et al.*, 2008) |
|  | ACD-5 | 4.87 (inhibitory)<br>6.48 (excitatory) | Li<sup>+</sup>> K<sup>+</sup>=Na<sup>+</sup> | IC<sub>50</sub>=131<br>μM | IC<sub>50</sub>=164 μM | intestine | (Kaulich *et al.*, 2021) |
|  | DEL-4 | 5.7 | Na<sup>+</sup>=Li<sup>+</sup>>K<sup>+</sup> | IC<sub>50</sub>=179<br>μM | IC<sub>50</sub>=12.25<br>μM | neurons | (Petratou *et al.*, submitted) |
|  | DELM-1 | 5.75 | Li<sup>+</sup>>Na<sup>+</sup>>K<sup>+</sup> | IC<sub>50</sub>=120<br>μM | EC<sub>50</sub>=238.3<br>μM | glia | (Han *et al.*, 2013) |
|  | UNC-105 | 6.42 | Na<sup>+</sup>=K<sup>+</sup> | IC<sub>50</sub>=<br>14.56 μM | IC<sub>50</sub>=31.94<br>μM | muscle | (Garcia-Anoveros *et al.*, 1998; Jospin & Allard, 2004) |
| acid- activated | ACD-2 | 5.03 | Li<sup>+</sup>=Na<sup>+</sup>>K<sup>+</sup> | IC<sub>50</sub>=84<br>μM | IC<sub>50</sub>=46.39<br>μM | head<br>neurons/glia |  |
|  | ASIC-1 | 4.49 | K<sup>+</sup>=Li<sup>+</sup>>Na<sup>+</sup> | IC<sub>50</sub>=134<br>μM | IC<sub>50</sub>=132.4<br>μM | neurons |  |
|  | DEL-9 | 4.27 | Na<sup>+</sup>=Li<sup>+</sup>=K<sup>+</sup> | *Acid-induced currents are insensitive* | IC<sub>50</sub>=58.33<br>μM | muscle,<br>neurons |  |

Table showing the channel properties described here, incorporated with those reported in the referenced indicated.

## Discussion

The acid-sensing members of the DEG/ENaCs, the ASICs, are proton receptors that can sense changes in extracellular pH in neuronal and non-neuronal tissues. *C. elegans* is a genetic model system that pioneered research into DEG/ENaCs, however, relatively little is known about pH-sensing by *C. elegans* DEG/ENaCs or their modulation by other molecules. We have shown here that there are at least three *C. elegans* acid-activated DEG/ENaCs, ASIC-1, ACD-2 and DEL-9. All three acid-activated channels are cation channels that are activated by increasing proton concentrations and inhibited by the trace element zinc. ASIC-1 and ACD-2 can also be blocked by the anti-hypertensive drug amiloride in a dose-dependent manner.

Our findings raise the question of the channels’ functional roles *in vivo*. In particular, it is unclear whether these channels would encounter a prolonged acidic environment under physiological conditions, especially as the acid-sensing DEG/ENaCs differ from the vertebrate ASICs in that they do not desensitise during the course of the proton stimulation (whereas the vertebrate ASICs do so within milliseconds). For instance, the murine ASIC1 is activated by protonergic neurotransmission, which might constitute a short increase in acidification due to neurotransmission, but could also represent a highly variable acidic environment depending on the rate of exocytosis (Du et al., 2014). Protons are co-packed in presynaptic vesicles with other neurotransmitters by the action of the proton pump Vacuolar-type ATPase (V-ATPase) (Gowrisankaran & Milosevic, 2020), and then co-released into the synaptic cleft, inducing a brief local drop in pH of 0.2 to 0.6 units (Miesenbock *et al*., 1998; Du *et al*., 2014; Zeng *et al*., 2015). This in turn stimulates postsynaptic receptors such as the ASICs (Soto *et al*., 2018). Similarly, presynaptic vesicles of glutamatergic hippocampal neurons also co-release millimolar concentrations of Zn^2+^ during synaptic transmission (Assaf & Chung, 1984; Howell *et al*., 1984; Frederickson & Moncrieff, 1994) which could in turn modulate ASIC channels during neurotransmission.

A role in neurotransmission may also be relevant to the *in vivo* function of *C. elegans* channels, in particular ASIC-1. We have shown that ASIC-1, like the murine ASIC1, can be activated by external protons in a concentration-dependent manner, suggesting that, ASIC-1 is involved in synaptic transmission in a similar way. This would fit well with behavioural and genetic evidence that it localizes to presynaptic terminals of dopaminergic neurons and enhances dopamine release required for associative learning (Voglis & Tavernarakis, 2008). Our electrophysiological characterisation of ASIC-1 *in vitro* supports the authors’ proposed working model in which ASIC-1 at the presynaptic terminal is activated by a local drop in pH during the release of dopamine from the pre-synaptic terminal, and this activation of ASIC-1 then promotes sustained dopaminergic signalling (Voglis & Tavernarakis, 2008). Expression in head neurons or glia suggests a likely role for ACD-2 in modulating synaptic function, similar to that described for the *C. elegans* ASIC-1, or modulating neuronal function as described for the glial DELM-1, DELM-2 or ACD-1 (Voglis & Tavernarakis, 2008; Wang *et al*., 2008; Han *et al*., 2013). DEL-9 is expressed in the GABAergic motor neuron AVL, which synapses on to the enteric muscle and regulates the expulsion step of the defecation motor program (McIntire *et al*., 1993), and the egg-laying muscles which are responsible for the expulsion of eggs. This suggests that, in common with the acid-inhibited channels ACD-5 and FLR-1/ACD-3/DEL-5 (Kaulich *et al*., 2021), it could function in the coordination of rhythmic behaviours.

We also expanded the characterisation of acid-inhibited channels, of which we identified four further members, ACD-5, DELM-1, DEL-4 and UNC-105, in addition to the previously described ACD-1 (Wang *et al*., 2008). This shows that even on exposure to the same stimulus (here protons), these channels show a remarkable diversity of functions, most likely responding to the demands in their particular environment (i.e. in the intestinal lumen (Kaulich *et al*., 2021), duct of the kidney (Collier & Snyder, 2009), and the synaptic cleft (Du *et al*., 2014)). UNC-105 functions in the body wall muscle and mutations that increase Na^+^ influx cause hypercontraction indicating a role in maintaining cell excitability (Park & Horvitz, 1986b; Garcia-Anoveros *et al*., 1998). Previous research, involving electrophysiological recording from muscle, has already suggested that *C. elegans* body muscles are proton-sensitive (Jospin *et al*., 2004): under voltage-clamp and current-clamp conditions, decreasing external pH from 7.2 to 6.1 led to a reversible depolarization of muscle cells (Jospin & Allard, 2004). However, in an *unc-105* null mutant the pH-sensitive current could still be observed, and acid-evoked depolarization is moreover suggestive of the involvement of an acid-activated, muscle-expressed channel, such as DEL-9 rather than an acid-inhibited channel like UNC-105. The roles of acid-sensing DEG/ENaCs in body muscle clearly merits further investigation.

Finally, DEG/ENaCs can also exert an effect on neuronal function from surrounding glia. Mutation of the *C. elegans* DEG/ENaC, *acd-1*, expressed in the amphid socket cells, exacerbates these sensory deficits, and deficits caused by mutations of genes implicated in sensory functions in other amphid neurons (Wang *et al*., 2008; Wang *et al*., 2012). Artificially increasing intracellular Ca^2+^ levels in one of these neurons bypassed the need for ACD-1, supporting the idea that ACD-1 modulates neuronal excitability (Wang *et al*., 2008; Wang *et al*., 2012). This idea is supported by the second example: DELM-1 (which we have shown is acid-inhibited, like ACD-1) and DELM-2 are expressed in the glia cells associated with nose touch neurons, the OLQs and IL1, on which they appear to exert a similar effect (Han *et al*., 2013). Vertebrate ASICs are also expressed in glia and, for example, some of the roles identified in learning may in fact be glia-based (Hill & Ben-Shahar, 2018), so disentangling glial from neuronal functions represents an exciting avenue for future investigations. In summary, *C. elegans* acid-sensitive ion channels appear to be involved in regulating the general excitability of a wide range of cell types, including muscle, glia, epithelia and neurons, and in a wide range of functional contexts. Precise channel localisation (for example, to identify localisation at specific synapses), and correlation with behavioural testing, are the necessary next steps for exploring *in vivo* function.

The expanded group of *C. elegans* DEG/ENaC thus encompasses a huge variety of channel properties, with respect to proton-dependence profiles, their interactions with amiloride, zinc and NSAIDs and mechanosensitivity (Chalfie & Sulston, 1981; Chatzigeorgiou *et al*., 2010; Geffeney *et al*., 2011; Fechner *et al*., 2021). These distinct functional capabilities do not necessarily cluster with overall sequence similarity. For example, in our phylogram (Figure 1), whereas ACD-1, ACD-5 and DELM-1 cluster closely to each other, the two other acid-inhibited subunits, DEL-4 and UNC-105, are closer to ASIC-1 and the mechanosensitive members, and none of the three acid-activated subunits cluster together. Disentangling the molecular basis of this diversity of function, and comparison across phyla, represents an important avenue for better understanding structure-function of the DEG/ENaCs. For instance, solving the structure of the chicken ASIC1 N-terminus and sequence comparison with other members revealed conserved residues important for ion-selectivity and gating (Yoder & Gouaux, 2020). Likewise, correlating sequence disparities with differences in function can play a key role in revealing their molecular basis.

Finally, our initial characterisation of these channels also provides a starting point for compound screens against these ASICs for drug discovery, both *in vivo* in the worm and in *Xenopus* oocytes, the former being particularly well-adapted to high throughput screening. Anti-helminthic drugs act on ion channels found in neurons and muscles and our characterisation has shown that *C. elegans* acid-activated DEG/ENaCs are present in both tissues, making them a good potential avenue with which to target parasitic relatives.

## Acknowledgements

We are very grateful to Kyuhyung Kim’s lab (Daegu Gyeongbuk Institute of Science and Technology) for providing us with their *C. elegans* DEG/ENaC transcriptional reporter plasmids and sharing their unpublished expression data with us. We thank members of the Schafer, Taylor and de Bono labs (MRC LMB), Beets and Timmerman (KU Leuven) and Pless (University of Copenhagen) labs, Iris Hardege, Vikram B. Kasaragod (MRC LMB), and Ewan St. John Smith (University of Cambridge) for helpful discussions. We are grateful to the LMB support facilities, in particular Ben Sutcliffe, Jonathan Howe, and Nick Barry from the Light Microscopy Team, and Sue Hubbard, Mark Cussens, and Martyn Howard and their team for preparing solutions and NGM plates. This work was supported by the Medical Research Council, as part of United Kingdom Research and Innovation (also known as UK Research and Innovation) [MRC file reference number MC-A023-5PB91] and by the Wellcome Trust [Grant reference number WT103784MA]. For the purpose of Open Access, the MRC Laboratory of Molecular Biology has applied a CC BY public copyright licence to any Author Accepted Manuscript (AAM) version arising from this submission.

## Notes

### Competing Interest Statement

The authors have declared no competing interest.

## References

Adams CM, Snyder PM & Welsh MJ. (1999). Paradoxical stimulation of a DEG/ENaC channel by amiloride. J Biol Chem 274, 15500–15504.

Assaf SY & Chung SH. (1984). Release of endogenous Zn2+ from brain tissue during activity. Nature 308, 734–736.

Awayda MS & Subramanyam M. (1998). Regulation of the epithelial Na+ channel by membrane tension. J Gen Physiol 112, 97–111.

Baconguis I, Bohlen CJ, Goehring A, Julius D & Gouaux E. (2014). X-ray structure of acid-sensing ion channel 1-snake toxin complex reveals open state of a Na(+)-selective channel. Cell 156, 717–729.

Baron A, Schaefer L, Lingueglia E, Champigny G & Lazdunski M. (2001). Zn2+ and H+ are coactivators of acid-sensing ion channels. J Biol Chem 276, 35361–35367.

Bentley PJ. (1968). Amiloride: a potent inhibitor of sodium transport across the toad bladder. J Physiol 195, 317–330.

Besson T, Lingueglia E & Salinas M. (2017). Pharmacological modulation of Acid-Sensing Ion Channels 1a and 3 by amiloride and 2-guanidine-4-methylquinazoline (GMQ). Neuropharmacology 125, 429–440.

Bhagatwala J, Harris RA, Parikh SJ, Zhu H, Huang Y, Kotak I, Seigler N, Pierce GL, Egan BM & Dong Y. (2014). Epithelial sodium channel inhibition by amiloride on blood pressure and cardiovascular disease risk in young prehypertensives. J Clin Hypertens (Greenwich) 16, 47–53.

Bianchi L, Gerstbrein B, Frokjaer-Jensen C, Royal DC, Mukherjee G, Royal MA, Xue J, Schafer WR & Driscoll M. (2004). The neurotoxic MEC-4(d) DEG/ENaC sodium channel conducts calcium: implications for necrosis initiation. Nat Neurosci 7, 1337–1344.

Boiko N, Kucher V, Stockand JD & Eaton BA. (2012). Pickpocket1 Is an Ionotropic Molecular Sensory Transducer. J Biol Chem 287, 39878–39886.

Canessa CM, Horisberger JD & Rossier BC. (1993). Epithelial sodium channel related to proteins involved in neurodegeneration. Nature 361, 467–470.

Canessa CM, Schild L, Buell G, Thorens B, Gautschi I, Horisberger JD & Rossier BC. (1994). Amiloride-sensitive epithelial Na+ channel is made of three homologous subunits. Nature 367, 463–467.

Cao J, Packer JS, Ramani V, Cusanovich DA, Huynh C, Daza R, Qiu X, Lee C, Furlan SN, Steemers FJ, Adey A, Waterston RH, Trapnell C & Shendure J. (2017). Comprehensive single-cell transcriptional profiling of a multicellular organism. Science 357, 661–667.

Carattino MD & Della Vecchia MC. (2012). Contribution of residues in second transmembrane domain of ASIC1a protein to ion selectivity. J Biol Chem 287, 12927–12934.

Chalfie M & Sulston J. (1981). Developmental genetics of the mechanosensory neurons of Caenorhabditis elegans. Dev Biol 82, 358–370.

Chalfie M & Wolinsky E. (1990). The identification and suppression of inherited neurodegeneration in Caenorhabditis elegans. Nature 345, 410–416.

Chatzigeorgiou M, Grundy L, Kindt KS, Lee WH, Driscoll M & Schafer WR. (2010). Spatial asymmetry in the mechanosensory phenotypes of the C. elegans DEG/ENaC gene mec-10. Journal of neurophysiology 104, 3334–3344.

Chen J, Winarski KL, Myerburg MM, Pitt BR & Sheng S. (2012). Probing the structural basis of Zn2+ regulation of the epithelial Na+ channel. J Biol Chem 287, 35589–35598.

Chen X, Polleichtner G, Kadurin I & Grunder S. (2007). Zebrafish acid-sensing ion channel (ASIC) 4, characterization of homo- and heteromeric channels, and identification of regions important for activation by H+. J Biol Chem 282, 30406–30413.

Ciccarelli FD, Doerks T, von Mering C, Creevey CJ, Snel B & Bork P. (2006). Toward automatic reconstruction of a highly resolved tree of life. Science 311, 1283–1287.

Collier DM & Snyder PM. (2009). Extracellular protons regulate human ENaC by modulating Na+ self-inhibition. J Biol Chem 284, 792–798.

De Stasio EA, Mueller KP, Bauer RJ, Hurlburt AJ, Bice SA, Scholtz SL, Phirke P, Sugiaman-Trapman D, Stinson LA, Olson HB, Vogel SL, Ek-Vazquez Z, Esemen Y, Korzynski J, Wolfe K, Arbuckle BN, Zhang H, Lombard-Knapp G, Piasecki BP & Swoboda P. (2018). An Expanded Role for the RFX Transcription Factor DAF-19, with Dual Functions in Ciliated and Nonciliated Neurons. Genetics 208, 1083–1097.

Driscoll M & Chalfie M. (1991). The mec-4 gene is a member of a family of Caenorhabditis elegans genes that can mutate to induce neuronal degeneration. Nature 349, 588–593.

Du J, Reznikov LR, Price MP, Zha XM, Lu Y, Moninger TO, Wemmie JA & Welsh MJ. (2014). Protons are a neurotransmitter that regulates synaptic plasticity in the lateral amygdala. Proc Natl Acad Sci U S A 111, 8961–8966.

Dulai JS, Smith ESJ & Rahman T. (2021). Acid-sensing ion channel 3: An analgesic target. Channels (Austin) 15, 94–127.

Elkhatib W, Smith CL & Senatore A. (2019). A Na(+) leak channel cloned from Trichoplax adhaerens extends extracellular pH and Ca(2+) sensing for the DEG/ENaC family close to the base of Metazoa. J Biol Chem 294, 16320–16336.

Fazia T, Pastorino R, Notartomaso S, Busceti C, Imbriglio T, Cannella M, Gentilini D, Morani G, Ticca A, Bitti P, Berzuini C, Dalmay T, Battaglia G & Bernardinelli L. (2019). Acid sensing ion channel 2: A new potential player in the pathophysiology of multiple sclerosis. Eur J Neurosci 49, 1233–1243.

Fechner S, D’Alessandro I, Wang L, Tower C, Tao L & Goodman MB. (2021). DEG/ENaC/ASIC channels vary in their sensitivity to anti-hypertensive and non-steroidal anti-inflammatory drugs. J Gen Physiol 153.

Frederickson CJ & Moncrieff DW. (1994). Zinc-containing neurons. Biol Signals 3, 127–139.

Friese MA, Craner MJ, Etzensperger R, Vergo S, Wemmie JA, Welsh MJ, Vincent A & Fugger L. (2007). Acid-sensing ion channel-1 contributes to axonal degeneration in autoimmune inflammation of the central nervous system. Nat Med 13, 1483–1489.

Garcia-Anoveros J, Garcia JA, Liu JD & Corey DP. (1998). The nematode degenerin UNC-105 forms ion channels that are activated by degeneration- or hypercontraction-causing mutations. Neuron 20, 1231–1241.

Geffeney SL, Cueva JG, Glauser DA, Doll JC, Lee TH, Montoya M, Karania S, Garakani AM, Pruitt BL & Goodman MB. (2011). DEG/ENaC but not TRP channels are the major mechanoelectrical transduction channels in a C. elegans nociceptor. Neuron 71, 845–857.

Gerlt JA. (2017). Genomic Enzymology: Web Tools for Leveraging Protein Family Sequence-Function Space and Genome Context to Discover Novel Functions. Biochemistry 56, 4293–4308.

Gerlt JA, Bouvier JT, Davidson DB, Imker HJ, Sadkhin B, Slater DR & Whalen KL. (2015). Enzyme Function Initiative-Enzyme Similarity Tool (EFI-EST): A web tool for generating protein sequence similarity networks. Biochim Biophys Acta 1854, 1019–1037.

Gowrisankaran S & Milosevic I. (2020). Regulation of synaptic vesicle acidification at the neuronal synapse. IUBMB Life 72, 568–576.

Han L, Wang Y, Sangaletti R, D’Urso G, Lu Y, Shaham S & Bianchi L. (2013). Two novel DEG/ENaC channel subunits expressed in glia are needed for nose-touch sensitivity in Caenorhabditis elegans. J Neurosci 33, 936–949.

Hardege I, Xu S, Gordon RD, Thompson AJ, Figg N, Stowasser M, Murrell-Lagnado R & O’Shaughnessy KM. (2015). Novel Insertion Mutation in KCNJ5 Channel Produces Constitutive Aldosterone Release From H295R Cells. Mol Endocrinol 29, 1522–1530.

Hill AS & Ben-Shahar Y. (2018). The synaptic action of Degenerin/Epithelial sodium channels. Channels (Austin, Tex) 12, 262–275.

Howell GA, Welch MG & Frederickson CJ. (1984). Stimulation-induced uptake and release of zinc in hippocampal slices. Nature 308, 736–738.

Huang M, Gu G, Ferguson EL & Chalfie M. (1995). A stomatin-like protein necessary for mechanosensation in C. elegans. Nature 378, 292–295.

Husson SJ, Costa WS, Wabnig S, Stirman JN, Watson JD, Spencer WC, Akerboom J, Looger LL, Treinin M, Miller DM, 3rd, Lu H & Gottschalk A. (2012). Optogenetic analysis of a nociceptor neuron and network reveals ion channels acting downstream of primary sensors. Curr Biol 22, 743–752.

Jiang Q, Zha XM & Chu XP. (2012). Inhibition of human acid-sensing ion channel 1b by zinc. Int J Physiol Pathophysiol Pharmacol 4, 84–93.

Jospin M & Allard B. (2004). An amiloride-sensitive H+-gated Na+ channel in Caenorhabditis elegans body wall muscle cells. J Physiol 559, 715–720.

Jospin M, Mariol MC, Segalat L & Allard B. (2004). Patch clamp study of the UNC-105 degenerin and its interaction with the LET-2 collagen in Caenorhabditis elegans muscle. The Journal of physiology 557, 379–388.

Kaminski J, Gibson MK, Franzosa EA, Segata N, Dantas G & Huttenhower C. (2015). High-Specificity Targeted Functional Profiling in Microbial Communities with ShortBRED. PLoS computational biology 11, e1004557–e1004557.

Katoh K, Misawa K, Kuma K & Miyata T. (2002). MAFFT: a novel method for rapid multiple sequence alignment based on fast Fourier transform. Nucleic Acids Res 30, 3059–3066.

Katoh K & Standley DM. (2013). MAFFT multiple sequence alignment software version 7: improvements in performance and usability. Mol Biol Evol 30, 772–780.

Kaulich E, Ackley BD, Tang Y, Hardege I, Schafer WR & Walker DS. (2021). Distinct roles for two Caenorhabditis elegans acid-sensing ion channels in an ultradian clock. bioRxiv.

Keiser J & Utzinger J. (2008). Efficacy of current drugs against soil-transmitted helminth infections: systematic review and meta-analysis. JAMA 299, 1937–1948.

Kellenberger S, Gautschi I & Schild L. (2003). Mutations in the epithelial Na+ channel ENaC outer pore disrupt amiloride block by increasing its dissociation rate. Mol Pharmacol 64, 848–856.

Kuraku S, Zmasek CM, Nishimura O & Katoh K. (2013). aLeaves facilitates on-demand exploration of metazoan gene family trees on MAFFT sequence alignment server with enhanced interactivity. Nucleic Acids Res 41, W22–28.

Larkin MA, Blackshields G, Brown NP, Chenna R, McGettigan PA, McWilliam H, Valentin F, Wallace IM, Wilm A, Lopez R, Thompson JD, Gibson TJ & Higgins DG. (2007). Clustal W and Clustal X version 2.0. Bioinformatics 23, 2947–2948.

Letunic I & Bork P. (2019). Interactive Tree Of Life (iTOL) v4: recent updates and new developments. Nucleic Acids Res 47, W256–W259.

Levin BJ, Huang YY, Peck SC, Wei Y, Martínez-Del Campo A, Marks JA, Franzosa EA, Huttenhower C & Balskus EP. (2017). A prominent glycyl radical enzyme in human gut microbiomes metabolizes trans-4-hydroxy-l-proline. Science (New York, NY) 355.

Li T, Yang Y & Canessa CM. (2009). Interaction of the aromatics Tyr-72/Trp-288 in the interface of the extracellular and transmembrane domains is essential for proton gating of acid-sensing ion channels. J Biol Chem 284, 4689–4694.

Li WG, Yu Y, Huang C, Cao H & Xu TL. (2011). Nonproton ligand sensing domain is required for paradoxical stimulation of acid-sensing ion channel 3 (ASIC3) channels by amiloride. J Biol Chem 286, 42635–42646.

Loytynoja A & Goldman N. (2010). webPRANK: a phylogeny-aware multiple sequence aligner with interactive alignment browser. BMC Bioinformatics 11, 579.

Matasic DS, Holland N, Gautam M, Gibbons DD, Kusama N, Harding AMS, Shah VS, Snyder PM & Benson CJ. (2021). Paradoxical Potentiation of Acid-Sensing Ion Channel 3 (ASIC3) by Amiloride via Multiple Mechanisms and Sites Within the Channel. Front Physiol 12, 750696.

McIntire SL, Jorgensen E, Kaplan J & Horvitz HR. (1993). The GABAergic nervous system of Caenorhabditis elegans. Nature 364, 337–341.

Miesenbock G, De Angelis DA & Rothman JE. (1998). Visualizing secretion and synaptic transmission with pH-sensitive green fluorescent proteins. Nature 394, 192–195.

O’Brodovich H, Canessa C, Ueda J, Rafii B, Rossier BC & Edelson J. (1993). Expression of the epithelial Na+ channel in the developing rat lung. Am J Physiol 265, C491–496.

Ortega-Ramirez A, Vega R & Soto E. (2017). Acid-Sensing Ion Channels as Potential Therapeutic Targets in Neurodegeneration and Neuroinflammation. Mediators Inflamm 2017, 3728096.

Palmer LG & Frindt G. (1986). Amiloride-sensitive Na channels from the apical membrane of the rat cortical collecting tubule. Proc Natl Acad Sci U S A 83, 2767–2770.

Park EC & Horvitz HR. (1986a). C. elegans unc-105 mutations affect muscle and are suppressed by other mutations that affect muscle. Genetics 113, 853–867.

Park EC & Horvitz HR. (1986b). Mutations with dominant effects on the behavior and morphology of the nematode Caenorhabditis elegans. Genetics 113, 821–852.

Paukert M, Babini E, Pusch M & Grunder S. (2004). Identification of the Ca2+ blocking site of acid-sensing ion channel (ASIC) 1: implications for channel gating. J Gen Physiol 124, 383–394.

Schild L, Schneeberger E, Gautschi I & Firsov D. (1997). Identification of amino acid residues in the alpha, beta, and gamma subunits of the epithelial sodium channel (ENaC) involved in amiloride block and ion permeation. J Gen Physiol 109, 15–26.

Schneider CA, Rasband WS & Eliceiri KW. (2012). NIH Image to ImageJ: 25 years of image analysis. Nat Methods 9, 671–675.

Shannon P, Markiel A, Ozier O, Baliga NS, Wang JT, Ramage D, Amin N, Schwikowski B & Ideker T. (2003). Cytoscape: a software environment for integrated models of biomolecular interaction networks. Genome Res 13, 2498–2504.

Soto E, Ortega-Ramirez A & Vega R. (2018). Protons as Messengers of Intercellular Communication in the Nervous System. Front Cell Neurosci 12.

Take-Uchi M, Kawakami M, Ishihara T, Amano T, Kondo K & Katsura I. (1998). An ion channel of the degenerin/epithelial sodium channel superfamily controls the defecation rhythm in Caenorhabditis elegans. Proc Natl Acad Sci U S A 95, 11775–11780.

Tan G, Muffato M, Ledergerber C, Herrero J, Goldman N, Gil M & Dessimoz C. (2015). Current Methods for Automated Filtering of Multiple Sequence Alignments Frequently Worsen Single-Gene Phylogenetic Inference. Syst Biol 64, 778–791.

Teiwes J & Toto RD. (2007). Epithelial sodium channel inhibition in cardiovascular disease. A potential role for amiloride. Am J Hypertens 20, 109–117.

Vina E, Parisi V, Cabo R, Laura R, Lopez-Velasco S, Lopez-Muniz A, Garcia-Suarez O, Germana A & Vega JA. (2013). Acid-sensing ion channels (ASICs) in the taste buds of adult zebrafish. Neurosci Lett 536, 35–40.

Voglis G & Tavernarakis N. (2008). A synaptic DEG/ENaC ion channel mediates learning in C. elegans by facilitating dopamine signalling. EMBO J 27, 3288–3299.

Voilley N. (2004). Acid-sensing ion channels (ASICs): new targets for the analgesic effects of non-steroid anti-inflammatory drugs (NSAIDs). Curr Drug Targets Inflamm Allergy 3, 71–79.

Voilley N, de Weille J, Mamet J & Lazdunski M. (2001). Nonsteroid Anti-Inflammatory Drugs Inhibit Both the Activity and the Inflammation-Induced Expression of Acid-Sensing Ion Channels in Nociceptors. J Neurosci 21, 8026–8033.

Vullo S & Kellenberger S. (2020). A molecular view of the function and pharmacology of acid-sensing ion channels. Pharmacol Res 154, 104166.

Waldmann R, Champigny G, Bassilana F, Heurteaux C & Lazdunski M. (1997). A proton-gated cation channel involved in acid-sensing. Nature 386, 173–177.

Waldmann R, Champigny G, Voilley N, Lauritzen I & Lazdunski M. (1996). The mammalian degenerin MDEG, an amiloride-sensitive cation channel activated by mutations causing neurodegeneration in Caenorhabditis elegans. The Journal of biological chemistry 271, 10433–10436.

Wang Y, Apicella A, Jr., Lee SK, Ezcurra M, Slone RD, Goldmit M, Schafer WR, Shaham S, Driscoll M & Bianchi L. (2008). A glial DEG/ENaC channel functions with neuronal channel DEG-1 to mediate specific sensory functions in C. elegans. EMBO J 27, 2388–2399.

Wang Y, D’Urso G & Bianchi L. (2012). Knockout of glial channel ACD-1 exacerbates sensory deficits in a C. elegans mutant by regulating calcium levels of sensory neurons. J Neurophysiol 107, 148–158.

Wang Y, Matthewman C, Han L, Miller T, Miller DM, 3rd & Bianchi L. (2013). Neurotoxic unc-8 mutants encode constitutively active DEG/ENaC channels that are blocked by divalent cations. J Gen Physiol 142, 157–169.

Wiemuth D & Grunder S. (2010). A single amino acid tunes Ca2+ inhibition of brain liver intestine Na+ channel (BLINaC). J Biol Chem 285, 30404–30410.

Williamson SM, Robertson AP, Brown L, Williams T, Woods DJ, Martin RJ, Sattelle DB & Wolstenholme AJ. (2009). The nicotinic acetylcholine receptors of the parasitic nematode Ascaris suum: formation of two distinct drug targets by varying the relative expression levels of two subunits. PLoS Pathog 5, e1000517.

Wolstenholme AJ & Rogers AT. (2005). Glutamate-gated chloride channels and the mode of action of the avermectin/milbemycin anthelmintics. Parasitology 131 Suppl, S85–95.

Yoder N & Gouaux E. (2020). The His-Gly motif of acid-sensing ion channels resides in a reentrant ‘loop’ implicated in gating and ion selectivity. Elife 9.

Zallot R, Oberg N & Gerlt JA. (2019). The EFI Web Resource for Genomic Enzymology Tools: Leveraging Protein, Genome, and Metagenome Databases to Discover Novel Enzymes and Metabolic Pathways. Biochemistry 58, 4169–4182.

Zallot R, Oberg NO & Gerlt JA. (2018). ‘Democratized’ genomic enzymology web tools for functional assignment. Curr Opin Chem Biol 47, 77–85.

Zeng WZ, Liu DS, Liu L, She L, Wu LJ & Xu TL. (2015). Activation of acid-sensing ion channels by localized proton transient reveals their role in proton signaling. Sci Rep 5, 14125.

Zhang P & Canessa CM. (2002). Single channel properties of rat acid-sensitive ion channel −1alpha, −2a, and −3 expressed in Xenopus oocytes. J Gen Physiol 120, 553–566.

