## Supplemental data for "Physiological insight into the conserved properties of *Caenorhabditis elegans* acid-sensing DEG/ENaCs"

**KEY RESOURCE TABLE**

**Bacterial strains, chemicals, peptides and recombinant proteins.**

| **REAGENT OR RESOURCE** | **SOURCE** | **IDENTIFIER** |
|---|---|---|
| **Bacterial strains** | | |
| *E. coli* OP50 | CGC | OP50 |

| **REAGENT OR RESOURCE** | **SOURCE** | **IDENTIFIER** |
|---|---|---|
| **Chemicals, Peptides and Recombinant proteins** | | |
| Amiloride hydyrochloride | Sigma-Aldrich | |
| Zinc chloride | Sigma-Aldrich | |
| Direct-zol RNA Miniprep | Zymo Research | |
| SuperScript™ II Reverse Transcriptase | Invitrogen™ | |
| pUCIDT-KAN-egas-1 cDNA | IDT | pEK308 |
| pUCIDT-KAN-egas-2 cDNA | IDT | pEK309 |
| pUCIDT-KAN-del-5_F9F3.4 | IDT | pEK155 |

***Strains used.***

| ***REAGENT OR RESOURCE*** | **SOURCE** | **IDENTIFIER** |
|---|---|---|
| ***Experimental Models: Organism and Strains*** | | |
| *C. elegans var. Bristol N2* | Caenorhabditis Genetics Center (CGC) | N2 (wild-type) |
| *ljEX1344 [Pacd-2::GFP (50ng/µl); Punc-122::GFP (50ng/µl)]* | Current paper | AQ4647 |
| *ljEx1361 [Pdel-9::GFP (50ng/µl); Punc-122::GFP (50ng/µl)]* | Current paper | AQ4672 |
| *ljEx1448 [Pasic-1::mKate2 (10ng/µl) ; Punc-122::GFP (50ng/µl)]* | Current paper | AQ4840 |

*Primers used for plasmid construction*

| **REAGENT OR RESOURCE** | | **SOURCE** | **IDENTIFIER** |
|---|---|---|---|
| **Primers for cloning promoters, gDNA and cDNA** | | | |
| KSM Hifiuni F | AGATCTGGTTACCACTAAACCAGCC | This paper | backbone for all KSM vectors if not stated otherwise |
| KSM hifiuni R | TGCAGGAATTCGATATCAAGCTTATCGATACC | | |
| acd-5 fragment_KSM_F | TTGGGCCCCTCGAGGTCGACATGCGACGCGTAAGAAACC | This paper | pEK171 |
| acd-5 fragment_KSM_R | CTCCATTCGGGTGTTCTTGATTATGCTTCATGTATCACAGCTGGC | | |
| KSM vector_acd-5_F | AGGTTTCTTACGCGTCGCATGTCGACCTCGAGGGGCC | | |
| KSM vector_acd-5_F | CTGTGATACATGAAGCATAATCAAGAACACCCGAATGGAGTCTCT | | |
| acd-2_KSM_F | TTGGGCCCCTCGAGGTCGACATGCATCTCGAGGACGGTC | This paper | pEK214 |
| acd-2_KSM_R | CTCCATTCGGGTGTTCTTGATTAACGAGGAGACAAGGATGATGGTAAAG | | |
| KSM_acd-2_R | GGACCGTCCTCGAGATGCATGTCGACCTCGAGGGGC | | |
| KSM_acd-2_F | CATCCTTGTCTCCTCGTTAATCAAGAACACCCGAATGGAGTCT | | |
| F28A12.1_ACD-4_KSM_F2 | TATCGAATTCCTGCAATGAATAGAAAACGAAAATTATCGTGCTTTGTATCTGTC | This paper | pEK252 |
| F28A12.1_ACD-4_KSM_R2 | GTGGTAACCAGATCTTCATAGTTTTATAATAATTCCCAGATC | | |
| Y69H2.2_egas-3_KSM_F1 | CTTGATATCGAATTCCTGCAATGATTTTCCTGCTTTTCCTCATATTCCC | This paper | pPM003 |
| Y69H2.2_egas-3_KSM_R1 | GTTTAGTGGTAACCAGATCTCTACTTTTCATATTTCTGGCACAAAACCATAAACA | | |
| Y69H2.11_egas-1_KSM_F | GTTTAGTGGTAACCAGATCTTCACTTTCCATACTTCTTACAACATAACGTGAATAG | This paper | pPM001 |
| Y69H2.11_egas-1_KSM_R | CTTGATATCGAATTCCTGCAATGCTACTATTCCTCTTCTTTTTCCCGG | | |

| | | | |
|---|---|---|---|
| Y69H2.12_egas-2_KSM_F<br>Y69H2.12_egas-2_KSM_R | CTTGATATCGAATTCCTGCAATGATTTTCCTGCTTTTCCTCATATTCCC<br>GTGGTAACCAGATCTTCACTTTCTACAACATATTGTCAAAACTCCGAAC | This paper | pPM002 |
| F55G1.13_egas-4_KSM_F<br>F55G1.13_egas-4_KSM_R | CTTGATATCGAATTCCTGCAATGTTGCTGCTATGGTTTTTTCTTCCG<br>GTTTAGTGGTAACCAGATCTTCAAAGACGTTTGTTGAACAAAAGTATGAC | This paper | pPM004 |
| T28B8.5_del-4_KSM_F<br>T28B8.5_del-4_KSM_R | CTTGATATCGAATTCCTGCAATGGGTGTATTTTGGACCGGC<br>GTTTAGTGGTAACCAGATCTTCAATCATTAGAATGAGGCTTTGGTGGAAC | This paper | pEK230 |
| F16F9.5_KSM_F<br>F16F9.5_mec-10_KSM_R | CGAATTCCTGCAgtacaaaattcaaaaaATGAATCGAAACCCGC<br>GTTTAGTGGTAACCAGATCTTCAATACTCATTTGCAGCATTTTCTC | This paper | pEK253 |
| C47C12.6_deg-1_KSM_R<br>C47C12.6_deg-1_KSM_F | GTGGTAACCAGATCTTTATATTGATACGAAAGCGTCTGACTTTCGCC<br>CGAATTCCTGCAGCCCGGGGGATCCACTAGTATGTCGAACCATCACAGTAAAAC | This paper | pEK251 |
| E02H4.1_del-1_KSM_F<br>E02H4.1_del-1_KSM_R | GATATCGAATTCCTGCAATGGCAAGGAAGTATATTGATATTTTAAAAAAATCAAAAATG<br>GTTTAGTGGTAACCAGATCTTCAATTATTATTTGTGGATACTCCTTTTTCCGCA | This paper | pEK329 |
| F59F3.4_del-5_KSM_F1<br>F59F3.4_del-5_KSM_R1 | CTTGATATCGAATTCCTGCAATGACGAGTGTCTCGTTTGGT<br>GTTTAGTGGTAACCAGATCTTTAAAAATCATTCATAGGCATATTTTTGGTGAATGCT | This paper | pEK228 |
| F_T28D9.7_del-10_KSM<br>R_T28D9.7_del-10_KSM | CTTGATATCGAATTCCTGCAATGGTCCGCATGGCTGAG<br>GTTTAGTGGTAACCAGATCTCTACACGTAAGAATGTTTATCATCATCCTCTTCG | This paper | pEK229 |
| R_del-9_C18B2.6_KSM<br>F_del-9_C18B2.6_KSM | GTTTAGTGGTAACCAGATCTTCATATGGGAGGCGTCGTTTCT<br>CTTGATATCGAATTCCTGCAATGTACATGAATGGAAATTTTCCCGAGAC | This paper | pEK219 |
| R13A1.4c_KSM_F<br>R13A1.4c_unc-8c_KSM_R1 | TCGAATTCCTGCATGATTCCAAAATATACATTTCCACGTCGC<br>GTGGTAACCAGATCTCTATTTGCTCATTAACTCCTTTGTTGATTCATTTG | This paper | pEK285 |
| F25D1.4_degt-1_F<br>F25D1.4_degt-1_R | CTTGATATCGAATTCCTGCAATGCCTCGAAAAAGAAGATCTGAAGAC<br>GTTTAGTGGTAACCAGATCTTTATATAAATTGTGGTTTTAGGAATATATTACTTTTCTTTCGTTCAC | This paper | pEK254 |

| | | | |
|---|---|---|---|
| F58G6.6a_del-2a_F<br>F58G6.6a_del-2a_R | CTTGATATCGAATTCCTGCAATGTTCTGCTTTCTGCAGTTACCG<br>GTTTAGTGGTAACCAGATCTTCACATATTGTCAGGCAAGTTTCTTCTGG | This paper | pEK247 |
| F58G6.6b_del-2b_F<br>F58G6.6b_del-2b_R | CTTGATATCGAATTCCTGCAATGAAAGGGCACACAGATTTTGATG<br>GTTTAGTGGTAACCAGATCTTCACATATTGTCAGGCAAGTTTCTTCTGG | This paper | pEK248 |
| asic-2_KSM_F<br>asic-2_KSM_R | CTTGATATCGAATTCCTGCAATGCGCGGTGGCG<br>GTTTAGTGGTAACCAGATCTTTATTTCTTCTTTTTCTCCTCATCTCCTTTATTCTCGA | This paper | pEK264 |
| C27C12.5b_KSM_F1<br>C27C12.5b_KSM_R1 | CCCTCGAGGTCGACGGATGACTGAAACTTCAAATTGCTCCAG<br>CTTAGAGACTCCATTCGGGTTTAGAAATCACAATTTCCGAGATACACAGAATTTCTTTT | This paper | pEK207 |
| T28B8.5_del-4_KSM_F<br>T28B8.5_del-4_KSM_R | CTTGATATCGAATTCCTGCAATGGGTGTATTTTGGACCGGC<br>GTTTAGTGGTAACCAGATCTTCAATCATTAGAATGAGGCTTTGGTGGAAC | This paper | pEK230 |
| ZK770.1_KSM_F<br>ZK770.1_KSM_R | TATCGAATTCCTATGGGAAAGAACAGCTTAAAACGGG<br>GTAACCAGATCTATCAATTATCAAGATTAAACCCGTCTTTGTTTAAATTATAATCAG | This paper | pEK234 |
| del-3_KSM_F<br>del-3_KSM_R | CTTGATATCGAATTCCTGCAATGTGGCTCCGAGGACTTTT<br>GTTTAGTGGTAACCAGATCTTTATGTGTCTCCTGAAGCTACATCTTGAC | This paper | pEK236 |
| del-7_KSM_F<br>del-7_KSM_R | CTTGATATCGAATTCCTGCAATGAATTGTAGCTGTGGTCATCAAACAG<br>GTTTAGTGGTAACCAGATCTTTATAGATCCATTTCGCGATTTTCTCGAA | This paper | pEK237 |
| C24G7.4_KSM_F<br>C24G7.4_KSM_R1 | TGATATCGAATTCCTGCAGCATGCATCTCGAGGACGGTC<br>TCCATTCGGGTGTTCTTGAGTTAACGAGGAGACAAGGATGATGGTAAAGAGG | This paper | pEK216 |
| F02D10.5_KSM_F1<br>F02D10.5_KSM_R1 | AGCTTGATATCGAATTCCTGATGGAAACGGAGACGGAAAGTG<br>TTCTTGAGGCTGGTTTAGTGTCAAATTAATTGTGATTTGAATATGGAGGATGTTGAAACT | This paper | pEK215 |
| ACD-1_KSM_F<br>ACD-1_KSM_R | GCTTGATATCGAATTCCTGCATGGAGCCAACTTTATCTCCAAATTATCGAAATG<br>GGTTTAGTGGTAACCAGATCTTAATATTTTTGAAAACTCGCGTTCCGGG | This paper | pEK216 |
| UNC-105_KSM_F<br><br>UNC-105_KSM_R<br>KSM_delm-1_F | GGTTTAGTGGTAACCAGATCTTTATGGTTCTTCTGGAGAGACTGGTCGATTACC<br><br>GCTTGATATCGAATTCCTGCAATGGAGAATGCGTCGTCAACTGCCC<br>CAAATTTACAAAAATATTAGTACCACTAAACCAGCCTCAAGAACACC | This paper<br><br><br>This paper | pEK385<br><br><br>pEK226 |

| | | | |
|---|---|---|---|
| KSM_delm-1_F | TTTGATCTGACCACTAAACCAGCCTCAAGAACACCC | | |
| C24G7.1_delm-2_KSM_F<br><br>C24G7.1_delm-2_KSM_R | GATATCGAATTCCTGCAGATGATTCCAACAATATCGAAGCCAAAAACC<br><br>GTGGTAACCAGATCTCAGATCAAAACGCTCCTTGATCCGAC | This paper | pEK384 |
| mec-4_KSM_F<br>mec-4_KSM_R | CTTGATATCGAATTCCTGCAATGTCATGGATGCAAAACCT<br>GTTTAGTGGTAACCAGATCTTCAGAAAGATCCAGACGCAATTTTCTTT | This paper | pEK266 |
| T21C9.3a_del-6a_F<br><br>T21C9.3a_del-6a_R | GTTTAGTGGTAACCAGATCTTTATTTCTTATTAACAGTCTTTTTATCAAAAGTATTTCCCTTCTTGAG<br><br>CTTGATATCGAATTCCTGCAATGGGTGCCAAGGTAAAGGAT | This paper<br><br>This paper | pEK249 |
| C11E4.3a_del-8_KSM_F1<br><br>C11E4.3a_del-8_KSM_F1 | CTTGATATCGAATTCCTGCAATGCCGGATAAAATCACAATTGCTG<br><br>GTTTAGTGGTAACCAGATCTTCACATGTTACTGTTGCCGAATAGTGG | This paper | pEK231 |

***Plasmids used.***

| **REAGENT OR RESOURCE** | **SOURCE** | **IDENTIFIER** |
|---|---|---|
| **Recombinant DNA (Plasmids)** | | |
| acd-5::KSM | (Kaulich et al., 2021) | pEK171 |
| Pacd-2::GFP | Kyuhyung Kim Lab | pEK186 |
| Pdel-9::GFP | Kyuhyung Kim Lab | pEK199 |
| acd-3::KSM | (Kaulich et al., 2021) | pEK207 |
| acd-2::KSM | current paper | pEK214 |
| flr-1::KSM | (Kaulich et al., 2021) | pEK215 |
| acd-1::KSM | current paper | pEK216 |
| del-9::KSM | current paper | pEK219 |
| del-5::KSM | (Kaulich et al., 2021) | pEK228 |
| del-10::KSM | current paper | pEK229 |

| | | |
|---|---|---|
| del-4::KSM | (Petratou *et al.*, submitted) | pEK230 |
| asic-1::KSM | current paper | pEK234 |
| del-3::KSM | current paper | pEK236 |
| del-7::KSM | current paper | pEK237 |
| Pasic-1 (3.5kb)::mKate2 | current paper | pEK240 |
| del-2a::KSM | current paper | pEK247 |
| del-2b::KSM | current paper | pEK248 |
| deg-1::KSM | current paper | pEK251 |
| acd-4::KSM | current paper | pEK252 |
| mec-10::KSM | current paper | pEK253 |
| degt-1::KSM | current paper | pEK254 |
| egas-1::KSM | current paper | pPM001 |
| egas-2::KSM | current paper | pPM002 |
| egas-3::KSM | current paper | pPM003 |
| egas-4::KSM | current paper | pPM004 |
| asic-2::KSM | current paper | pEK264 |
| unc-8 (isoform c)::KSM | current paper | pEK285 |
| del-1::KSM | current paper | pEK329 |
| unc-105(isoform h)::KSM | current paper | pEK385 |
| delm-1::KSM | current paper | pEK226 |
| delm-2::KSM | current paper | pEK384 |
| mec-4::KSM | current paper | pEK266 |
| del-6 (isoform a)::KSM | current paper | pEK249 |
| del-8 (isoform b)::KSM | current paper | pEK231 |

***Software, Algorithms and other equipment.***

| **Software and Algorithms** | |
|---|---|
| Graphpad | GraphPad Software inc |
| Roboocyte2+ | Multichannel Systems inc. |

| | |
|---|---|
| SnapGene | Insightful Science |
| GUIDANCE2 | (Sela et al., 2015) |
| MAFFT | (Katoh et al., 2002, Katoh and Standley, 2013) |
| iTOL | (Ciccarelli et al., 2006, Letunic and Bork, 2019, Kuraku et al., 2013) |
| EFI-EST | Enzyme Function Initiative (efi.igb.illinois.edu/) |

| **Other** | |
|---|---|
| Roboocyte | Multichannel Systems inc. |
| Roboinject | Multichannel Systems inc. |

**Supplementary figures**

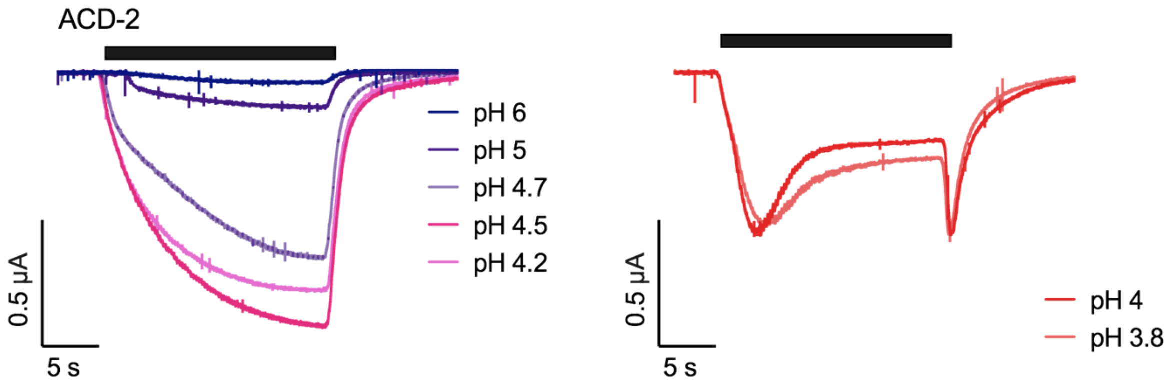


***Supplementary Figure 1: ACD-2 currents above and below pH 4.***
Representative traces from ACD-2 expressing oocytes (N = 6). Left: perfusion of pH 6 – pH 4.2, show a decrease in current (see main text Figure 5) while perfusion with <pH 4.2 resulted in a partial desensitisation of the channel. Black bars represent the time of perfusion with low pH from a neutral pH 7.4.

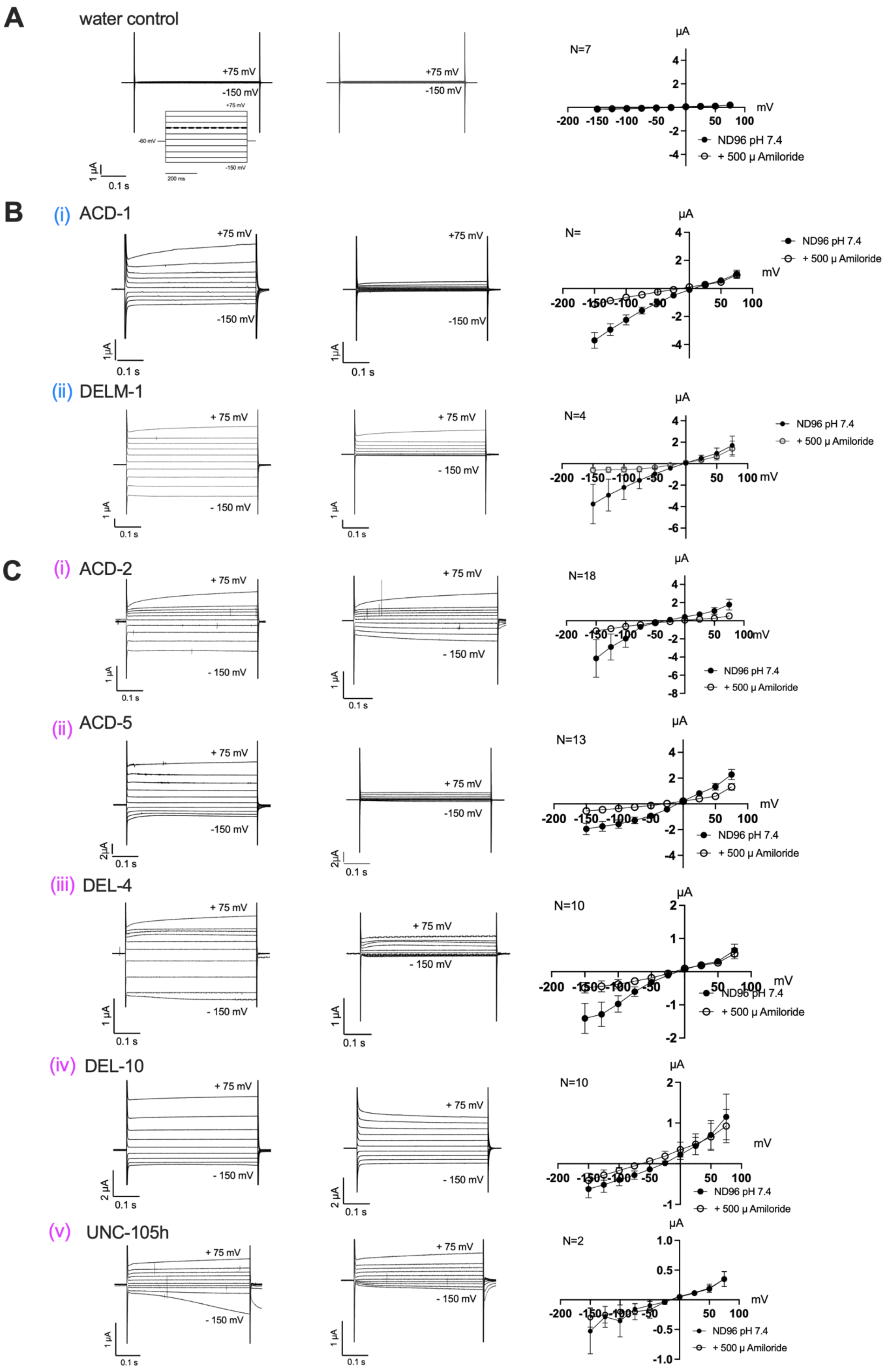
A
water control
+75 mV
-150 mV
N=7
ND96 pH 7.4
+ 500 μ Amiloride
B
(i) ACD-1
(ii) DELM-1
N=4
C
(i) ACD-2
N=18
(ii) ACD-5
N=13
(iii) DEL-4
N=10
(iv) DEL-10
N=10
(v) UNC-105h
N=2
μA
mV
0.1 s
1 μA

33

***Supplementary figure 2:* Amiloride can block currents of DEG/ENaC constructs expressing Xenopus oocytes.**

Representative transient currents and current–voltage (IV) relationships in the absence (left) and presence of 500 µM amiloride (middle) at pH 7.4. (A) Nuclease-free water-injected oocytes (negative control) are unaffected by amiloride. (B) Amiloride block. ACD-1 and DELM-1 (positive controls), ACD-2, ACD-5, DEL-4, DEL-10 and UNC-105h transient currents can be blocked by amiloride. *Xenopus* oocytes are perfused with a basal solution (ND96) only (filled circles), and in presence of the DEG/ENaC channel blocker amiloride (open circles). The oocyte membrane was clamped at −60 mV and voltage steps from -150 mV to +75 mV were applied as indicated. Currents are actual currents in µA (y-axis), voltage steps in mV (x-axis) as indicated. Error bars represent Mean ± SEM.

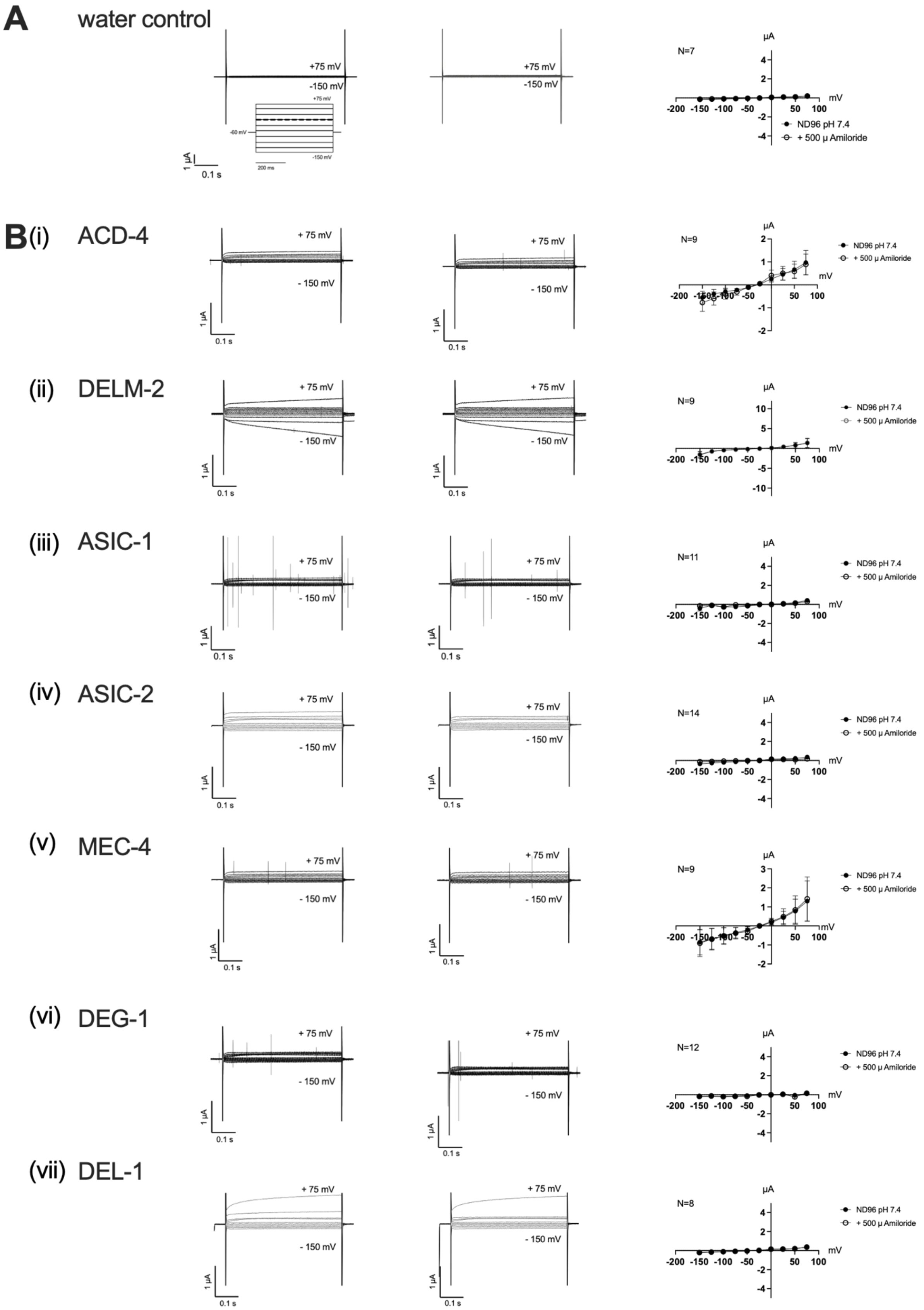
A
water control
+75 mV
-150 mV
1 μA
0.1 s
N=7
ND96 pH 7.4
+ 500 μ Amiloride
B(i)
ACD-4
N=9
(ii)
DELM-2
(iii)
ASIC-1
N=11
(iv)
ASIC-2
N=14
(v)
MEC-4
(vi)
DEG-1
N=12
(vii)
DEL-1
N=8
μA
mV

***Supplementary figure 3:* Amiloride-insensitive currents of Xenopus oocytes expressing DEG/ENaC constructs.**

First and second column: Representative transient currents and current–voltage (IV) relationships in the absence (left) and presence of 500 µM amiloride (middle) at pH 7.4. Third Colum: Averaged IV in the absence and presence of 500 µM amiloride at pH 7.4. (A) Nuclease-free water-injected oocytes (negative control) are unaffected by amiloride. ACD-4, DELM-2, ASIC-1, ASIC-2, MEC-4, DEG-1 and DEL-1 transients are unaffected by amiloride (FLR-1 and ACD-3 are published elsewhere (Kaulich et al., 2021)). Xenopus oocytes are perfused with a basal solution (ND96) only (filled circles), and in presence of the DEG/ENaC channel blocker amiloride (open circles). The oocyte membrane was clamped at −60 mV and voltage steps from -150 mV to +75 mV were applied as indicated. Currents are actual currents in µA (y-axis), voltage steps in mV (x-axis) as indicated. Error bars represent Mean ± SEM.

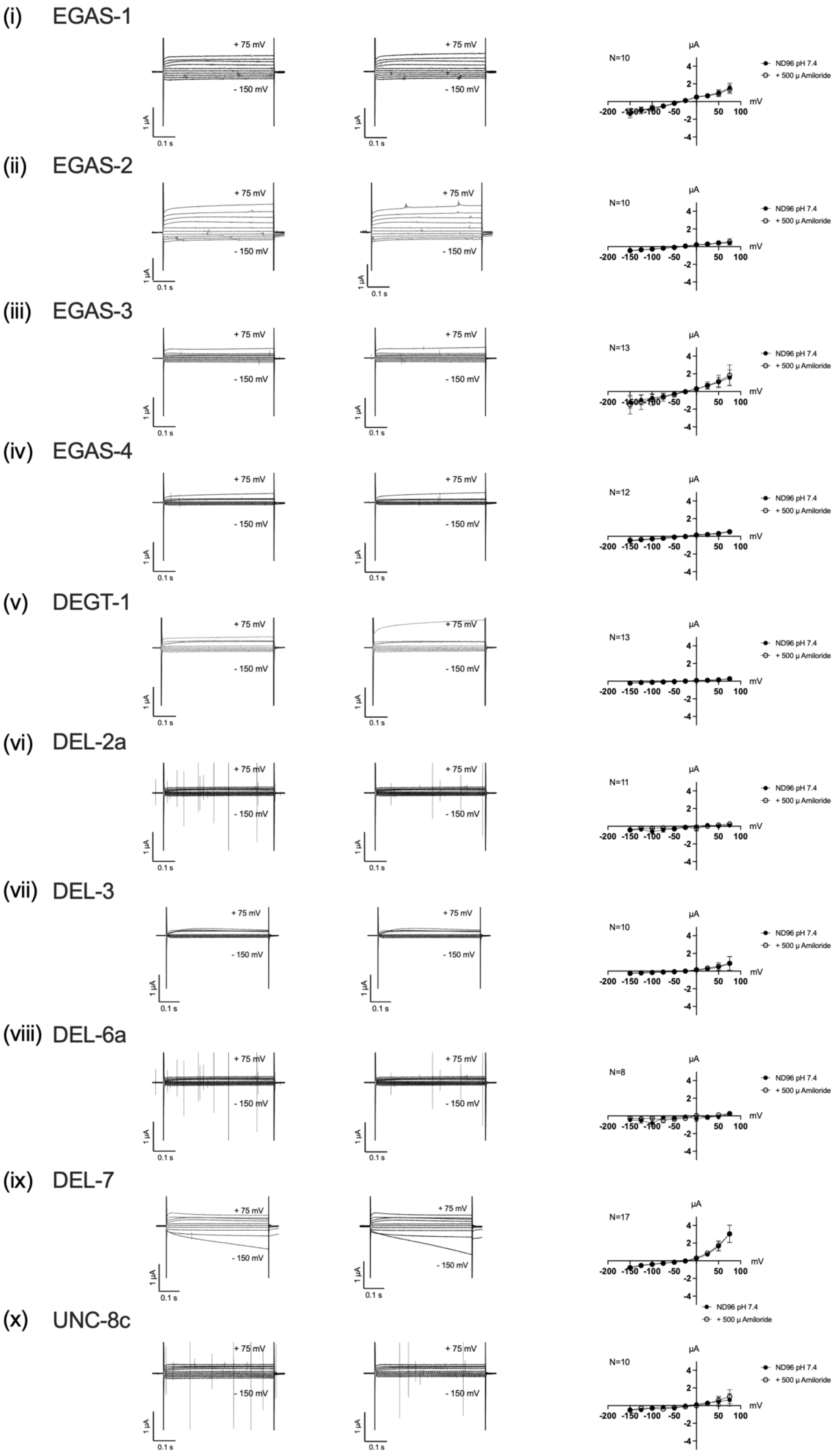
(i) EGAS-1
(ii) EGAS-2
(iii) EGAS-3
(iv) EGAS-4
(v) DEGT-1
(vi) DEL-2a
(vii) DEL-3
(viii) DEL-6a
(ix) DEL-7
(x) UNC-8c
+ 75 mV
- 150 mV
1 μA
0.1 s
N=10
N=10
N=13
N=12
N=13
N=11
N=10
N=8
N=17
N=10
μA
mV
ND96 pH 7.4
+ 500 μ Amiloride

***Supplementary figure 4:* Amiloride-insensitive currents of Xenopus oocytes expressing DEG/ENaC constructs.**

First and second column: Representative transient currents and current–voltage (IV) relationships in the absence (left) and presence of 500 µM amiloride (middle) at pH 7.4. Third Colum: Averaged IV in the absence and presence of 500 µM amiloride at pH 7.4 Nuclease-free water-injected oocytes (negative control) are unaffected by amiloride. EGAS-1, EGAS-2, EGAS-3, EGAS-4, DEGT-1, DEL-2, DEL-3, DEL6a, DEL-7 and UNC-8 transients are unaffected by amiloride. *Xenopus* oocytes are perfused with a basal solution (ND96) only (filled circles), and in presence of the DEG/ENaC channel blocker amiloride (open circles). The oocyte membrane was clamped at −60 mV and voltage steps from -150 mV to +75 mV were applied as indicated. Currents are actual currents in µA (y-axis), voltage steps in mV (x-axis) as indicated. Error bars represent Mean ± SEM.

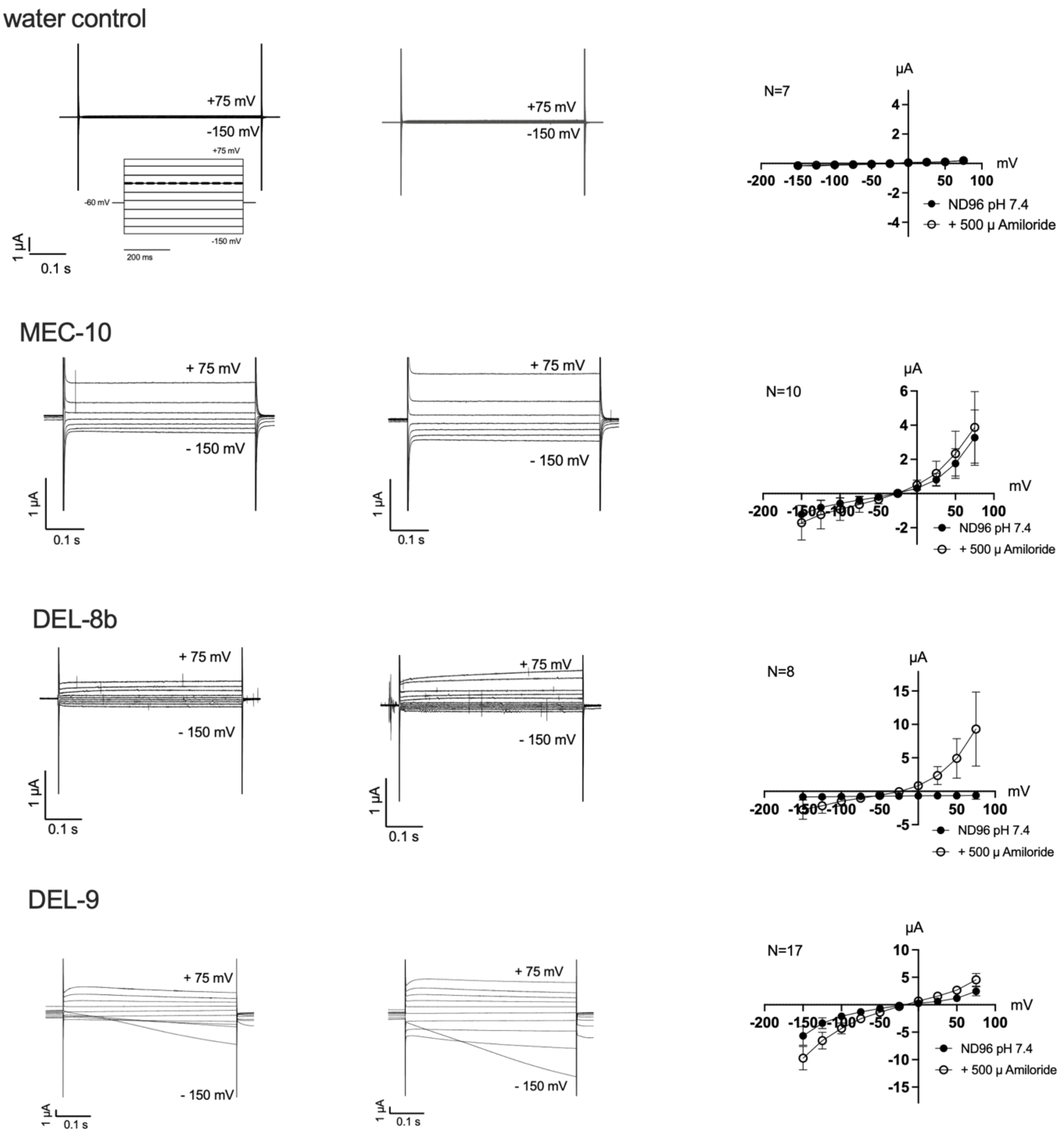


***Supplementary figure 5:* Amiloride can enhance currents of DEG/ENaC constructs expressing *Xenopus* oocytes.**

Representative transient currents and current–voltage (IV) relationships in the absence (left) and presence of 500 µM amiloride (middle) at pH 7.4. (A) Nuclease-free water-injected oocytes (negative control) are unaffected by amiloride. Amiloride current-potentiation. MEC-10, DEL-5, DEL-8b and DEL-9 transient currents are enhanced in the presence of amiloride. DEL-5 amiloride-activated transients are published elsewhere (Kaulich et al., 2021). *Xenopus* oocytes are perfused with a basal solution (ND96) only (filled circles), and in presence of the DEG/ENaC channel blocker amiloride (open circles). The oocyte membrane was clamped at −60 mV and voltage steps from -150 mV to +75 mV were applied as indicated. Currents are actual currents in µA (y-axis), voltage steps in mV (x-axis) as indicated. Error bars represent Mean ± SEM.

*Supplementary Table 1: DEG/ENaC Accession numbers.*

| ***Organism*** | ***Protein name (accession number)*** |
|---|---|
| ***Caenorhabditis elegans*** | FLR-1 (UniProtKB/TrEMBL ID G5EGI5); ACD-1 (P91102); ACD-2 P91100); ACD-3 (G3MU02); ACD-4 (Q22970); ACD-5 (O01664); DELM-1 (O45402); DELM-2 (P91103); ASIC-1 (K7H9J0); ASIC-2 (Q22851); MEC-4 (P24612); DEL-4 (P91835); UNC-105 (Q09274); DEG-1 (P24585); DEL-1 (Q19038); MEC-10 (P34886); EGAS-1 (Q9U1T9); EGAS-2 (Q9U1T8); EGAS-3 (Q9XTS9); EGAS-4 (Q20852); DEGT-1 (Q19777); DEL-2 (G5ECD8); DEL-3 (Q93597); DEL-5 (G5EFH3); DEL-6 (Q8MPW0); DEL-7 (Q18651); DEL-8 (Q93205); UNC-8 (Q21974); DEL-9 (Q18077); DEL-10 (Q10025). |
| ***Drosophila melanogaster*** | PPK1 (Q7KT94); PPK2 (O46342); PPK3 (Q8MLR6); PPK4 (O61365); PPK5 (Q7KTW2); PPK6 (Q86LH3); PPK7 (Q9VME9); PPK8 (B7Z123); PPK9 (Q9W2B5); PPK10 (Q86LH1); PPK11 (Q9VL84); PPK12 (Q9W250); PPK13 (Q86LG9); PPK14 (Q86LG8); PPK15 (Q9VBF6); PPK16 (Q86LG7); PPK17 (Q9VJI4); PPK18 (Q9VL88); PPK19 (Q9VAJ3); PPK20 (Q86LG5); PPK21 (Q86LG4); PPK22 (Q8IMV2); PPK23 (Q9VX46); PPK24 (Q9V9Y5); PPK25 (A1Z6S4); PPK26 (Q9VS73); PPK27 (Q9VZN1); PPK28 (Q86LG1); PPK29 (A8DYP2); PPK30 (Q9VAJ5); PPK31 (A8JPJ8) |
| ***Rattus norvegicus*** | rASIC1 (P55926); rASIC2 (Q62962); rASIC3 (O35240); rASIC4 (Q9JHS6); rASIC5 (Q9R0W5); ENaCα (P37089); ENaCβ (P37090); ENaCγ (P37091) |
| ***Danio rerio*** | zAISC1A (Q708S7); zAISC1B (Q708S8); zAISC1C (Q708S6); zAISC2 (Q708S5); zAISC4A (Q708S4); zAISC4B (Q708S3) |
| ***Hydra vulgaris*** | HyNaC2 (A8DZR6); HyNaC3 (A8DZR7); HyNaC4 (A8DZR8); HyNaC5 (D3UD58); HyNaC6 (A0A0A0MP54); HyNaC7 (A0A0A0MP73); HyNaC8 (A0A0A0MP55); HyNaC9 (A0A0A0MP48); HyNaC10 (A0A0A0MP61); HyNaC11 (A0A0A0MP67); HyNaC12 (A0A0A0MP74) |
| ***Aplysia kurodai*** | FaNaC (Q4H3X6) |
| ***Platynereis dumerilii*** | pENaC4 (A0A2S1B6I2); pENaC7 (A0A2S1B6Q3); pENaC6 (A0A2S1B6R1); pMGIC (A0A2S1B6I3) |

| | |
|---|---|
| ***Lottia gigantean*** | LgFaNaC (V4C2H5) |
| ***Lymnaea stagnalis*** | LsFaNaC (Q9BJD0) |
| ***Helix aspersa*** | HaFaNaC (Q25011). |
| ***Helisoma trivolvis*** | HtFaNaC (Q9NBC7) |
| ***Trichoplax adhaerens*** | TadNaC1 (B3S0Z3); TadNaC2 (A0A5J6BSS6); TadNaC3 (A0A5J6BTF2); TadNaC4 (A0A5J6BSQ9); TadNaC5 (A0A5J6BSM6); TadNaC6 (A0A5J6BVG3); TadNaC7 (A0A5J6BSR6); TadNaC8 (A0A5J6BV03); TadNaC9 (A0A5J6BWR3); TadNaC10 (A0A5J6BSU1). |

### References


CICCARELLI, F. D., DOERKS, T., VON MERING, C., CREEVEY, C. J., SNEL, B. & BORK, P. 2006. Toward automatic reconstruction of a highly resolved tree of life. *Science,* 311**,** 1283-7.

KATOH, K., MISAWA, K., KUMA, K. & MIYATA, T. 2002. MAFFT: a novel method for rapid multiple sequence alignment based on fast Fourier transform. *Nucleic Acids Res,* 30**,** 3059-66.

KATOH, K. & STANDLEY, D. M. 2013. MAFFT multiple sequence alignment software version 7: improvements in performance and usability. *Mol Biol Evol,* 30**,** 772-80.

KAULICH, E., ACKLEY, B. D., TANG, Y., HARDEGE, I., SCHAFER, W. R. & WALKER, D. S. 2021. Distinct roles for two Caenorhabditis elegans acid-sensing ion channels in an ultradian clock. *bioRxiv*.

KURAKU, S., ZMASEK, C. M., NISHIMURA, O. & KATOH, K. 2013. aLeaves facilitates on-demand exploration of metazoan gene family trees on MAFFT sequence alignment server with enhanced interactivity. *Nucleic Acids Res,* 41**,** W22-8.

LETUNIC, I. & BORK, P. 2019. Interactive Tree Of Life (iTOL) v4: recent updates and new developments. *Nucleic Acids Res,* 47**,** W256-W259.

SELA, I., ASHKENAZY, H., KATOH, K. & PUPKO, T. 2015. GUIDANCE2: accurate detection of unreliable alignment regions accounting for the uncertainty of multiple parameters. *Nucleic Acids Res,* 43**,** W7-14.
